# Efficient Hi-C inversion facilitates chromatin folding mechanism discovery and structure prediction

**DOI:** 10.1101/2023.03.17.533194

**Authors:** Greg Schuette, Xinqiang Ding, Bin Zhang

## Abstract

Genome-wide chromosome conformation capture (Hi-C) experiments have revealed many structural features of chromatin across multiple length scales. Further understanding genome organization requires relating these discoveries to the mechanisms that establish chromatin structures and reconstructing these structures in three dimensions, but both objectives are difficult to achieve with existing algorithms that are often computationally expensive. To alleviate this challenge, we present an algorithm that efficiently converts Hi-C data into contact energies, which measure the interaction strength between genomic loci brought into proximity. Contact energies are local quantities unaffected by the topological constraints that correlate Hi-C contact probabilities. Thus, extracting contact energies from Hi-C contact probabilities distills the biologically unique information contained in the data. We show that contact energies reveal the location of chromatin loop anchors, support a phase separation mechanism for genome compartmentalization, and parameterize polymer simulations that predict three-dimensional chromatin structures. Therefore, we anticipate that contact energy extraction will unleash the full potential of Hi-C data and that our inversion algorithm will facilitate the widespread adoption of contact energy analysis.

**Significance Statement:** The three-dimensional organization of the genome is essential to many DNA-templated processes, and numerous experimental techniques have been introduced to characterize its features. High-throughput chromosome conformation capture experiments, or Hi-C, have proven particularly useful, reporting the interaction frequency between pairs of DNA segments *in vivo* and genome-wide. However, the polymer topology of chromosomes complicates Hi-C data analysis, which often employs sophisticated algorithms without explicitly accounting for the disparate processes affecting each interaction frequency. In contrast, we introduce a computational framework based on polymer physics arguments that efficiently removes the correlation between Hi-C interaction frequencies and quantifies how each local interaction influences genome folding globally. This framework facilitates the identification of mechanistically important interactions and the prediction of three-dimensional genome structures.

## Introduction

The three-dimensional organization of the genome is an essential component of the processes that regulate gene expression and encode cellular diversity [1–4]. This biological importance has motivated the development of many experimental techniques to characterize genome structures and understand the mechanisms leading to their formation. Hi-C experiments [5, 6] have proven particularly insightful, as the high-resolution and genome-wide information they report has revealed structural features including chromatin loops [7], topologically associating domains [8, 9], and A/B compartments [6].

Hi-C reports the interaction frequency between each pair of genomic loci *in vivo*, and the experimental output is typically presented in two-dimensional contact probability maps whose axes index the genetic sequence [10, 11]. This data contains the information needed to identify non-random contacts of functional significance, including the promoter-enhancer pairs involved with gene regulation [7, 10]. However, interpreting Hi-C data can be complicated, as the polymer structure of chromosomes introduces several features that obscure the functionally relevant information [10–12]. Notably, configuration entropy penalizes the interaction between genomic loci far apart in sequence and couples contact formation between distinct genomic pairs [5, 12–16]. Existing approaches often address the genomic distance effect by normalizing contact probabilities to the average value of all contact pairs having the same genomic separation [11, 17–19], but they rarely account for the correlation between contact probabilities [14, 15] that obscures the location of specific interactions [20–23]. Therefore, it remains difficult to determine whether a genomic pair exhibiting high contact probability is energetically favorable due to a specific interaction between them (driving) or entropically favorable due to topological constraints associated with a driving interaction occurring elsewhere (passive). Differentiating between driving and passive interactions is necessary to identify which are biologically significant, deepen mechanistic insight, and improve upon existing approaches to predict three-dimensional genome structures using Hi-C [16, 20, 21, 24–35].

The information theoretic approach introduced by Zhang and Wolynes decouples the contribution of specific interactions from topological constraints in Hi-C data by converting contact probabilities into interaction energies that measure the attraction between genomic loci brought into proximity [20, 21]. Their algorithm performs this task by inverting a polymer model of chromatin derived to reproduce Hi-C data while obeying the maximum entropy principle [36]. The resulting interaction energies are independent of one another because the energy function of the model explicitly separates the effects of local interactions and polymer topology, which couples contact formation. Therefore, interaction energies directly correspond to the biologically important, specific interactions that cause the unique features in Hi-C probability maps to form, providing direct insight to the forces that drive genome folding.

While the interpretation of interaction energies is biologically meaningful, extracting them from Hi-C contact probabilities is nontrivial. The existing solutions [20–23, 37, 38] to this problem require many time-consuming molecular dynamics (MD) simulations to be performed sequentially, so the algorithms suffer from high computational complexity. Additionally, interaction energies extracted by these approaches suffer from statistical noise because adequately sampling conformations of long polymers during each algorithmic step is computationally infeasible using MD simulations. Computationally efficient alternative approaches are highly desirable [39].

In this study, we introduce a new method to address the challenges associated with converting Hi-C probabilities into interaction energies. This new approach utilizes a contact-space representation of polymers, in which a set of binary variables represents polymer conformations by indicating which monomers are in contact. An approachable, Ising-like Hamiltonian accounts for the topology-driven coupling between contacts and the distance effect in this space, and we introduce analytical expressions that parameterize the Hamiltonian using contact statistics. Extending this model to chromatin systems, another analytical expression converts Hi-C contact probabilities into contact energies, which are the contact-space analogue of interaction energies with an identical interpretation. Testing this extraction algorithm with an *in silico* model showed that the extracted contact energies accurately identify the mechanistically important contacts. Furthermore, we found that contact energies extracted from micro-C data [40] of H1-hESC and HFFc6 cells successfully reveal the location of anchors in chromatin loops. Contact energies extracted from compartment-scale data support a phase-separation model, as interactions between loci in the same compartment are almost universally more favorable than interactions between loci in different compartments. Finally, we show that contact energies can parameterize polymer models that reasonably reproduce Hi-C data at different resolutions. We believe these results indicate that the algorithm introduced herein will greatly simplify the interpretation of Hi-C data and facilitate the prediction of genome structures using computer simulations.

## Methods

### Theory I: Maximum entropy inversion algorithm

As mentioned in the Introduction, the polymer structure of chromosomes couples the formation of interactions between distinct pairs of genomic loci [14, 15], so the contact probability between two genomic loci is influenced by specific interactions that occur both locally and at distant interaction sites. Accordingly, Hi-C contact probabilities obscure the location of many biologically significant interactions. Interaction energies, in contrast, directly measure the strength of these specific interactions, so they are unaffected by the effects of polymer topology. In this way, converting from contact probabilities to interaction energies distills the information needed to identify structurally significant interactions.

This conversion can be performed by inverting a polymer model of chromatin to reproduce Hi-C contact probabilities while obeying the maximum entropy principle [20, 21]. For example, starting with a homopolymer model that has the energy function *U*_hp_(***r***), the least-biased model that reproduces Hi-C data can be derived by maximizing the excess entropy as [41–45]

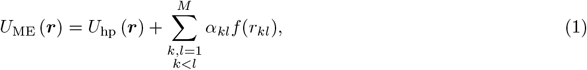

where *k* and *l* index all *M* monomers, ***r*** contains their Cartesian coordinates, *f* is a threshold function that determines whether two monomers are in contact given their Euclidean distance *r*_*kl*_, and *α*_*kl*_ is an interaction energy such that the unique set ***α*** ≡ {*α*_*kl*_} can be determined from experimental contact probabilities. Physically, ***α*** captures the influence of specific interactions by quantifying the interaction strength between each pair of genomic loci, whereas *U*_hp_(***r***) captures the polymer topological effects. Thus, inverting this model separates the physical phenomena that contribute to Hi-C contact probabilities, and ***α*** readily identifies structurally significant interactions.

Although this approach is theoretically sound, existing inversions [20–23, 37, 38] solve ***α*** by iteratively performing polymer simulations, which are computationally expensive and suffer from statistical noise. Thus, further developing robust and efficient inversions would expand the number of feasible use cases for interaction energies and simplify Hi-C data analysis.

### Theory II: Modeling chromatin and extracting energies in contact space

Rather than inverting the polymer model in three dimensions, we approach the problem in a binary contact space. The polymer is mapped from its Cartesian representation ***r*** to its binary representation ***q*** via the transformation

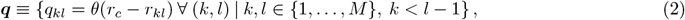

where *q*_*kl*_ = 1 (*q*_*kl*_ = 0) indicates that monomer pair (*k, l*) is (not) in contact. In place of the general threshold function *f*, this conversion determines contact with the Heaviside step function *θ* and contact distance *r*_c_, meaning *q*_*kl*_ is equal to

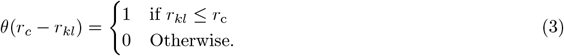

Imposing *k* < l − 1 ensures that each *q*_*kl*_ is unique while also ignoring nearest neighbors, which are trivially in contact. Proceeding, we index contact-space objects with *i, j* ∈ { (*k, l*) | *k, l* ∈ {1, …, *M*}, *k < l* − 1} to emphasize that they reside in a different mathematical space than objects indexed with *k* and *l*.

To capture topological effects in contact space, we approximate *U*_hp_(***r***) with the Ising-like expression

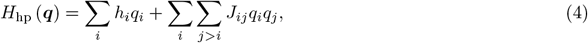

where *i* and *j* index the *N* = (*M* − 1)(*M* − 2)*/*2 unique contact pairs in ***q***. This Hamiltonian has found success in studies of protein folding [46, 47], and the linear and quadratic model parameters account for distinct effects of polymer topology contributing to contact probabilities [48]. In particular, *h*_*i*_ ∈ ***h*** ≡ {*h*_*i*_} increases with the sequence separation between the monomers involved with contact *i*, while the quadratic parameters *J*_*ij*_ ∈ ***J*** ≡ {*J*_*ij*_} capture the correlation between contact pairs.

Extending *H*_hp_ to a chromatin model that recreates Hi-C probabilities while maximizing information entropy, we find

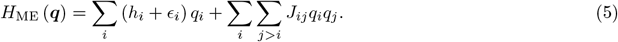

In analogy to *α*_*kl*_ ∈ ***α***, *ϵ*_*i*_ represents a *contact energy*, and ***ϵ*** ≡ {*ϵ*_*i*_} captures all system-specific, biologically interesting contributions to Hi-C probabilities. As such, though contact energies are a first-order interaction term much like *h*_*i*_, they capture the direct impact of pairwise attractions, while the *h*_*i*_ continue to capture topological effects. Conveniently, the ensemble average ⟨*q*_*i*_⟩_ME_ and the Hi-C contact probability *p*_*i*_ are equal, where the subscript in ⟨·⟩_A_ denotes the Hamiltonian used for the average. The simple form of the Hamiltonian allows ***ϵ*** to be extracted from Hi-C contact probabilities ***p*** ≡ {*p*_*i*_} with the following expression:

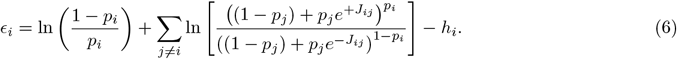

The Supporting Material details our derivation of Eq. 6, which relies on the inverse equations we developed to determine the topological parameters ***h*** and ***J***.

### Theory III: Determining the topological parameters of contact-space models

In order to solve the contact energies ***ϵ*** using Eq. 6, we must first determine the values of the topological parameters ***h*** and ***J***. Their exact solution is not known, but the parameters in a generalized Ising model can often be well-approximated with inversions that use equilibrium statistics or a set of configurations. These inversions have been widely used to analyze protein sequence statistics and infer which amino acid pairs are in direct contact [49, 50]. For homopolymers, however, we found that existing approaches, including the mean-field [51, 52] and optimization-based [53–56], provide unphysical (Fig. S1) and noisy (Fig. S2) results, respectively. To overcome these obstacles, we developed inverse equations well-suited to homopolymer models in contact space.

The Supporting Material provides our derivation of the inverse equation we used to compute the coupling matrix ***J***,

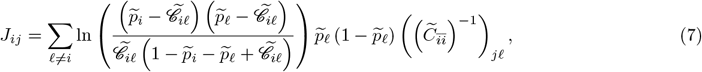

where 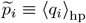 is the contact probability of contact *i*, 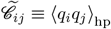 is the *connected correlation* between computed with contacts *i* and *j*, 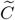 is the *N* × *N* covariance matrix such that 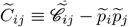, and 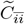 is the (*N* − 1) × (*N* − 1) submatrix of 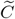 in which the *i*^th^ row and column have been removed. Similarly, the field parameters can be computed with

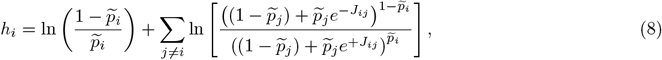

whose derivation is also introduced in the Supporting Material. Therefore, both ***h*** and ***J*** can be determined using the first-and second-order contact statistics of a homopolymer, which we approximated with polymer simulations.

### Theory IV: An uncoupled approximation provides a benchmark

The inversion algorithm defined in Eq. 6 rests on our insight that the coupling parameters of an Ising-like model can capture polymer topological effects well enough to provide sufficiently uncorrelated contact energies. To validate this concept, we will compare the results of our algorithm to those of its *uncoupled approximation*. This refers to the limit in which all *J*_*ij*_ = 0 in the contact-space model, which therefore treats the formation of each contact as an independent process. This reduces the homopolymer Hamiltonian to

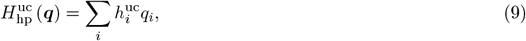

where the superscript ‘uc’ denotes objects specific to the uncoupled approximation. Because Eq. 9 contains only first-order model parameters, 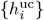 must account for the combined influence of all topological constraints. This remains true after extending the uncoupled model to the chromatin system, so Eq. 5 reduces to

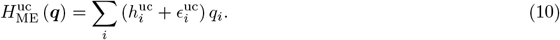

In this case, the sum in Eq. 8 equals 0, so Eq. 6 reduces to

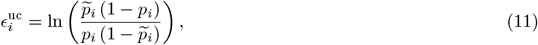

which compares the average free energy change associated with forming contact *i* in the chromatin model to that in the homopolymer. We note that, in the small-probability limit, 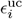 reduces to the “statistical potential” energies described by Shin, Shi, and Thirumalai [39], i.e.

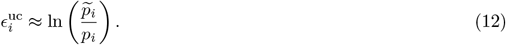

While not explicitly analyzed herein, this limit also represents the the normalization used to address the sequence separation-related probability decay in other methods, e.g. [17–19].

### Simulating the homopolymer to obtain contact statistics

We carried out MD simulations in reduced units, using the software LAMMPS [57] to obtain an ensemble of homopolymer configurations that provided contact statistics. The homopolymer consists of 500 monomers, and nearest-neighbor monomers are connected by class2 bonds, implemented as

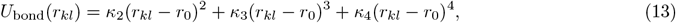

with bond length *r*_0_ = 1 and force constants *κ*_2_ = *κ*_3_ = *κ*_4_ = 20. Excluded volume effects were not included to facilitate the relaxation of polymer configurations. Langevin dynamics maintained the system temperature of 1.0 using a damping coefficient of 10. One billion total time steps of length Δ*t* = 0.005 were performed, and configurations were saved every 5000 time steps to provide 200,000 total configurations.

We mapped each simulated conformation to contact space by implementing Eq. 2 in Python with contact distance *r*_c_ = 1.76. The distance between monomers, *r*_*kl*_, was computed with mdtraj 1.9.6 [58], then the simulated contact probabilities and covariances were computed with PyTorch 1.11.0 [59]. All further analysis used NumPy 1.19.2 [60].

We computed statistics for contacts in the central 300-monomer region of the homopolymer. The statistics of the central region of a much longer polymer better approximate those of an infinite homopolymer, as this avoids edge effects. In addition, identical topological conditions affect many contacts and contact pairs in an infinite homopolymer, so they maintain identical contact probabilities and correlations, respectively [14, 15]. Accordingly, we replaced each contact statistic with the average of all values associated with topologically identical interactions; this increased the effective number of configurations contributing to the contact statistics, thereby minimizing statistical uncertainty in the simulated data. The Supporting Material describes this process in detail.

### Simulating polymers with specific interactions

In addition to the above homopolymer simulation, we performed simulations to generate *in silico* contact probability maps and to evaluate the quality of contact energies extracted from Hi-C data. These simulations again featured a polymer model consisting of 500 monomers connected by class2 bonds. We also applied non-bonded contact potentials between monomers *k* and *l* in the following form

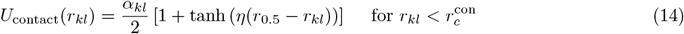

using decay rate *η* = 5, interaction distance *r*_0.5_ = 1.75, cutoff distance 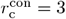. Our analysis focused on the central 200 monomers in the polymer to minimize edge effects, so we indexed the monomers with *k, l* ∈ {−149, …, 350} to maintain consistency between sections.

When using the extracted contact energies to recreate Hi-C probability maps, we also applied a Lennard-Jones potential between all monomer pairs to account for the excluded volume effect of chromatin and better match the separation dependence of contact probabilities observed in chromatin. The potential is defined as

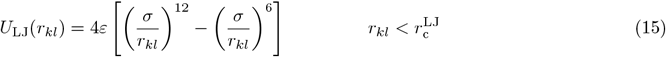

with *ε* = 0.35, *σ* = 1.0, and cutoff 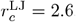. In addition, we reduced simulation length to 120,000,000 time steps when recreating Hi-C maps, performed 10 simulations in parallel, and increased save frequency to once per 1000 time steps. This provided 6,000,000 conformations to analyze while improving computation time.

### Preparing Hi-C data

Micro-C experiments, a Hi-C variant, provided the *in vivo* contact probabilities used throughout this work. The .mcool files containing the Micro-C dataset for hESC and HFF cells were downloaded from the 4DN Data Portal with accession numbers 4DNFI9GMP2J8 and 4DNFI9FVHJZQ, respectively [40]. We loaded balanced Hi-C contacts probabilities from these files at the resolution of interest for the entire chromosome of interest using Cooler 0.8.11 [61]. The average of all real-valued probabilities from contacts with the same loop size, *K* ≡ |*l* − *k*|, served as the unnormalized, background-averaged contact probability 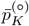. We further processed the 200-bin region of interest on its own, using cooltools 0.5.1 [62] to remove zero-valued probabilities via adaptive coarse-graining then undefined probabilities via linear interpolation. Finally, we divided all contact probabilities by 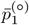 to normalize the data, satisfying our assumption that nearest neighbors have contact probability 1. The normalized data are denoted 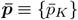 and ***p*** ≡ {*p*_*i*_} for loop size-averaged and full Hi-C contact probabilities, respectively.

### Obtaining compartment profiles

To aid the analysis of compartment-level contact energies, we obtained compartment profiles using the cooltools 0.5.1 [62] implementation of eigenvector decomposition [63] in Python. Positive and negative values in the first eigenvector, E1, correspond to genomic regions in the A- and B-compartment, respectively. In addition to the aforementioned Micro-C datasets [40], this process required the GC content associated with each genomic bin in the Hi-C data. To obtain this information, we downloaded the UCSC version of the December 2013 assembly of the human genome (GRCh38 Genome Reference Consortium accession number GCA 000001405.15) [64] in FASTA format from the UCSC Genome Browser database [65], loaded it using bioframe 0.3.3 [66], and determined the fraction of GC content in each genomic bin using the bioframe function frac gc. Finally, we used the eigs cis function in cooltools to perform eigenvector decomposition for the region of interest only.

### Preparing CTCF ChIP-seq signals

To support the relationship between prominent negative contact energies and loop-stabilizing interactions in chromatin systems, we plotted ChIP-seq data along the axes of contact energy maps extracted from *in vivo* Hi-C contact probabilities to highlight the genomic loci most strongly associated with CTCF. We downloaded preprocessed signal p-values in bigWig format from the ENCODE data portal [67] using accession numbers ENCFF147GRN [68] and ENCFF209TQB [69] for H1-hESC and HFFc6 cell types, respectively. Dividing the region of interest into nonoverlapping segments containing 100 base pairs each, we loaded the average signal p-value for each segment using pyBigWig 0.3.22 [70]. Each plot of ChIP-seq data is independently normalized to the largest signal in that plot.

## Results and Discussion

In the Methods section, we introduced an efficient algorithm that uses Hi-C contact probabilities to infer the strength of non-bonding contact potentials between each pair of monomers in polymer models of chromatin. This required representing polymers in contact space, where a set of binary variables, ***q*** ≡ {*q*_*i*_}, indicates which pairs of monomers are in contact and *i, j* ∈ {(*k, l*)} index all non-nearest-neighbor monomer pairs. The Ising-like Hamiltonian in Eq. 4 approximates the free energy function of a homopolymer in this space and, therefore, captures the generic effects of polymer topology. Specifically, ***h*** ≡ {*h*_*i*_} accounts for the entropic penalty associated with forming each contact, while ***J*** ≡ {*J*_*ij*_} combines the higher-order effects of polymer connectivity to capture contact correlations. Following a maximum entropy argument, Eq. 5 incorporates contact energies, *ϵ*_*i*_ ∈ ***ϵ***, to reproduce Hi-C contact probabilities and account for the local interactions between monomer pairs. The strength of these local interactions is independent of the polymer topological effects that couple contact formation and correlate Hi-C contact probabilities, so contact energies directly relate pairs of genomic loci to the biologically significant interactions that drive chromatin folding. Therefore, extracting contact energies from Hi-C contact probabilities distills the biologically significant information contained in Hi-C data and should simplify its analysis.

### Contact-space parameters capture polymer topological effects

The Ising-like Hamiltonian of homopolymers in contact space accounts for the effects of polymer topology on contact probability and the coupling between contact sites with ***h*** and ***J***, respectively. We computed these parameters using contact statistics obtained from MD simulations of a homopolymer, thus ensuring that topological effects alone contribute to their value. The Methods section provides simulation details, and Fig. 1 presents the results. As expected, the average contact probabilities, 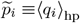, decrease with distance from the diagonal in Fig. 1*B* and loop size in Fig. 1*C* since larger loops are less likely to form. For illustrative purposes, we also plotted the correlation between a specific contact, that involving monomers 89 and 111, and other contacts in the upper triangle of Fig. 1*D*. However, the qualitative trends are not affected by this particular choice (see Fig. S3).

**Figure 1.**
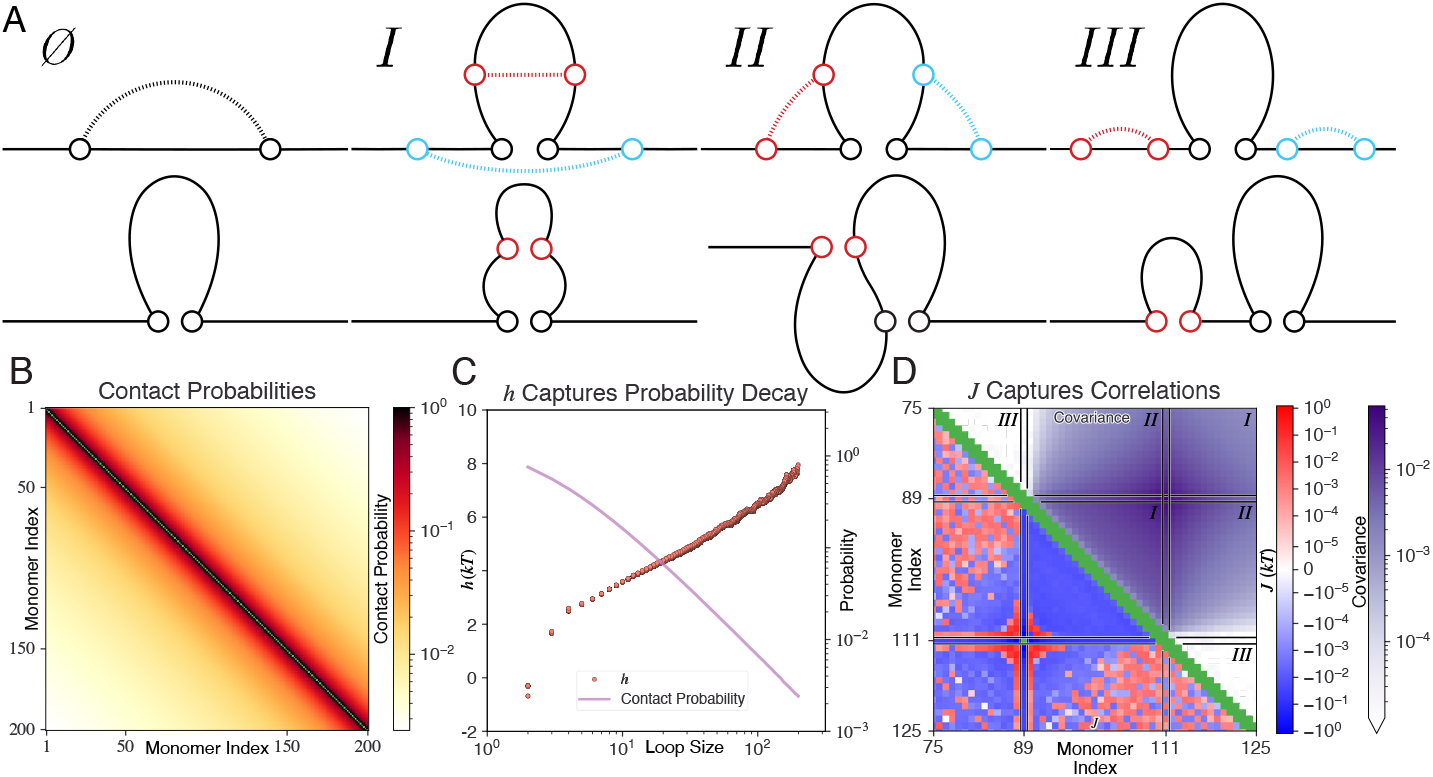
Parameters in the Ising-like Hamiltonian capture the effects of polymer topology in contact space. (A) Illustrating the relative position of monomers in various contact pairs (top) and the polymer structure associated with the formation of those contacts (bottom). Forming one contact yields a loop (∅). Presuming that the black contact exists, forming a second contact (red or blue) yields a structure belonging to one of three topological sets: One loop encloses the other in regime *I*; the loops partially overlap in regime *II*; and the loops are independent in regime *III*. While the polymer structure associated with the red contact is shown in each second-order diagram, forming the blue contact instead would yield a homeomorphic structure. (B) Contact probabilities between pairs of monomers calculated from MD simulations of a homopolymer model. (C) Dependence of the field parameters ***h*** (left axis) and average contact probabilities (right axis) as a function of the loop size. (D) The correlation (upper triangle) and *J*_*ij*_ coupling parameters (lower triangle) between contacts *i* = (89, 111) and *j* = (*k, l*) are compared. Black lines indicate the border between regimes *I, II*, and *III*, as defined in the main text and illustrated in (A). The discontinuous subsets of each regime correspond to the red and blue loops in (A). Green indicates nonexistent values.

Fig. 1*A* illustrates the topological considerations that give rise to the observed trends in these contact statistics. For example, distant monomers must be brought into proximity to form a loop (∅), which reduces the number of configurations the polymer segment lying between these monomers can assume. The proportion of configurations involving contact between two monomers decreases as their linear separation increases, so contact probability decreases as loop size increases. Similarly, the correlation between two contacts depends on the number of configurations available to the polymer when those contacts coexist. In particular, each pair of contacts belongs to one of three regimes representing distinct topologies [14, 15]. The right three panels of Fig. 1*A* illustrate the relative orientation of the monomers involved with contact pairs in each regime (top) and their associated topology (bottom). At one extreme, regime *I* contains all contact pairs in which one loop fully encloses the other. Forming either contact reduces the configurational entropy penalty associated with subsequently forming the other, so these contacts are positively correlated. At the other extreme, regime *III* contains all contact pairs with non-overlapping loops. In this case, the formation of one contact does not alter the configurational entropy penalty affecting the other, so the contacts are uncorrelated. Regime *II* represents the transition between these extremes, as it contains all contact pairs whose loops partially overlap. For otherwise-identical contact pairs, greater overlap between their loops increases their correlation.

We used the first- and second-order contact statistics, i.e. all 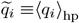 and 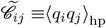, to compute ***J*** and ***h*** with Eqs. 7 and 8, respectively. Fig. 1*C* shows that contact probability and loop size are linearly related on a log scale, as anticipated [13–15]. *h*_*i*_ accounts for this effect by increasing with loop size. Meanwhile, the coupling parameters, *J*_*ij*_, track the correlation between contacts. As with pairwise correlations, we plotted coupling parameters associated with the contact between monomers 89 and 111 in the lower triangle of Fig. 1*D*. Most of the *J*_*ij*_ are negative in regimes *I* and *II*, supporting the correlation of the corresponding loops. The values are much smaller in magnitude in regime *III* due to the independence of the loops.

### An *in silico* model validates the energy extraction algorithm

With the Ising-like Hamiltonian of homopolymers parameterized, we extended the contact-space model to a chromatin-like model by introducing a set of contact energies, ***ϵ*** ≡ {*ϵ*_*i*_}, in the Hamiltonian in Eq. 5. This model can exactly reproduce Hi-C contact probability maps, and each map is associated with a unique set of contact energies. Our algorithm extracts this set of energies from Hi-C contact probabilities by applying the inverse Eq. 6 to all contacts. The resulting energies are local quantities because ***h*** ≡ {*h*_*i*_} and ***J*** ≡ {*J*_*ij*_} separately account for the effects of polymer topology, so they lack the cross-correlation that obscures biologically important information in Hi-C contact probabilities.

We assessed the quality of extracted contact energies by investigating an *in silico* chromatin model consisting of 200 monomers. In addition to the homopolymer interactions considered in the prior subsection, this model included a contact potential to promote loop formation. For simplicity, we only applied one non-bonded interaction, and it used the potential defined in Eq. 14 with *α*_*kl*_ = −4*k*_*B*_*T* between monomers 89 and 111. Conceptually, this represents a biologically significant interaction stabilizing a chromatin loop, e.g. a promoter-enhancer interaction. Fig. 2*A* portrays this step.

**Figure 2.**
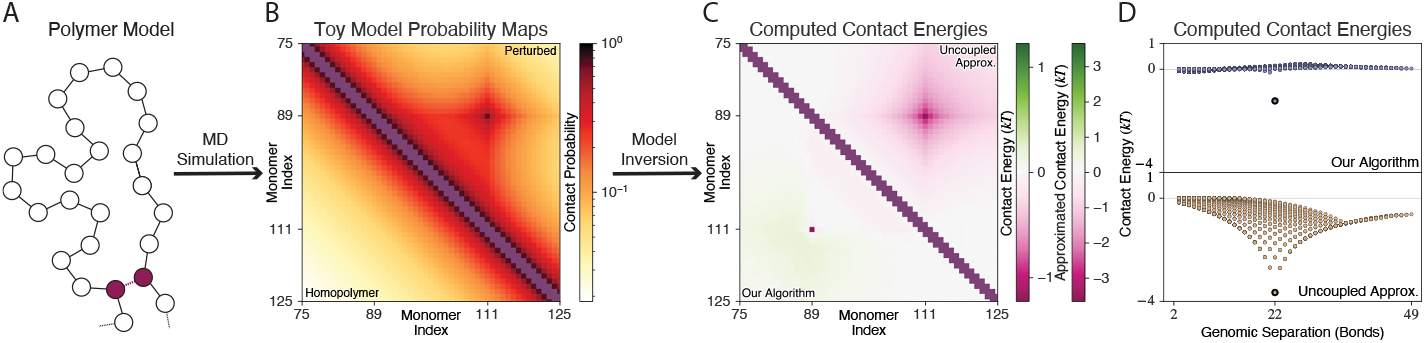
The energy extraction algorithm successfully identifies the specific contact of an *in silico* chromatin model. A flowchart illustrates our proposed algorithm using data from the *in silico* experiment. (A) Starting with the initial homopolymer model, we applied an attraction between monomers 89 and 111, illustrated in purple. (B) Comparing the simulated contact probabilities for this model (upper triangle) to those of the homopolymer model (lower triangle) reveals the effects of loop formation. (C) Comparing the contact energies extracted from the simulated contact probabilities using the coupled (***ϵ***, lower triangle) and uncoupled (***ϵ***^uc^, upper triangle) models illustrates the effectiveness of ***J*** in accounting for coupling in the polymer system. To best visualize its prominence, the loop-stabilizing interaction’s contact energy defines the minimum value of the colormap in each extraction approach (upper and lower triangle). (D) Plotting contact energies against their corresponding separation further emphasizes the uncorrelated nature of contact energies extracted using ***J***. Bold borders indicate the contact energy between monomers 89 and 111.

Polymer simulations of this model provided the *in silico* Hi-C contact probabilities, which Fig. 2*B* compares to the homopolymer probabilities. As anticipated, the favorable interaction between monomers 89 and 111 directly increased the contact probability at that site and indirectly increased the contact probability at many mechanistically insignificant contact sites. With no pairwise interaction between the monomers at these alternative sites, the coupling effect of polymer topology is singularly responsible for the change in their interaction frequency.

Next, we extracted contact energies using Eq. 6, where *p*_*i*_ represents the *in silico* Hi-C contact probabilities. As shown in the lower triangle of Fig. 2*C* and the upper plot of Fig. 2*D*, a prominent, negative-valued contact energy is associated with the attraction between monomers 89 and 111. In excellent agreement with the simulated model, near-zero contact energies appear at all other sites.

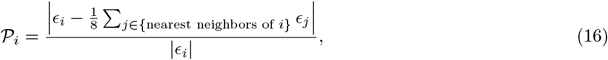

For comparison, the upper triangle of Fig. 2*C* and lower plot of Fig. 2*D* show the uncoupled approximation of these contact energies, meaning all *J*_*ij*_ = 0 when computing contact energies (Eq. 11). This approximation assumes all the contacts are independent, ignoring the coupling effect of polymer topology. A simple ratio quantifies this difference as which compares the difference between the contact energy of the stabilizing interaction and the average contact energies of its 8 nearest neighbor in the 2D plot. This yields 0.993 and 0.315 for the coupled and uncoupled contact energies, respectively, which implies that the mechanistically important interaction has a far more prominent contact energy when energies are extracted using our algorithm and the uncoupled approximation. The contact energies extracted for *in silico* models having multiple attractors display this same advantage (Fig. S4, Table S1). This crucial difference emphasizes the importance of ***J*** when extracting contact energies and illustrates the quality of *J*_*ij*_ computed with our inversion.

### Contact energies facilitate the identification of mechanistically important interactions

Chromatin looping and gene regulation are intricately connected, as persistent loops often bring promoter and enhancer regions into proximity [71–73]. Analytical techniques that use Hi-C data have expanded our understanding of the regulation of chromatin loops themselves by identifying the location of loop anchors and relating them to sequence information. However, identifying these interactions often requires sophisticated algorithms [7, 74–79], as the aforementioned influence of polymer topology complicates the relationship between the functional importance of an interaction and its probability. Because our algorithm separates these effects from the influence of local interactions, we anticipate that contact energy extraction will facilitate the identification of functionally important interactions.

To demonstrate their usefulness in this task, we extracted contact energies from Hi-C contact probabilities covering several regions of HFF and hESC cells; these were preprocessed according to the procedure described in the Methods section. In addition to extracting ***ϵ*** from the Hi-C contact probabilities ***p***, we extracted 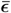 from the loop size-averaged contact probabilities 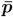. While the former represents the contact energies required to reproduce Hi-C probabilities, the latter represents the contact energies required to reproduce the probability decay of the relevant chromosome, meaning they capture nonspecific effects. Therefore, we analyzed the difference 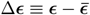 to further isolate the biologically interesting pairwise interactions. For comparison, we extracted contact energies using the uncoupled approximation and analyzed 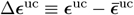.

Fig. 3*A* shows the contact energy map for a 1 Mb region of chromosome 1 in hESC cells at 5 kb resolution, with Δ***ϵ*** and Δ***ϵ***^uc^ shown in lower and upper triangle, respectively. For illustrative purposes, this figure includes the CTCF ChIP-seq signal along each axis. The correlation between contact energies in the uncoupled approximation is particularly evident in the loop regions bounded by each box. In contrast, the contact energies are much more localized when extracted with our algorithm, which predicts only a small minority of contacts to be strongly attractive. Importantly, these attractive contacts localize at CTCF peaks, supporting the role of these molecules in chromatin loop formation [80, 81].

**Figure 3.**
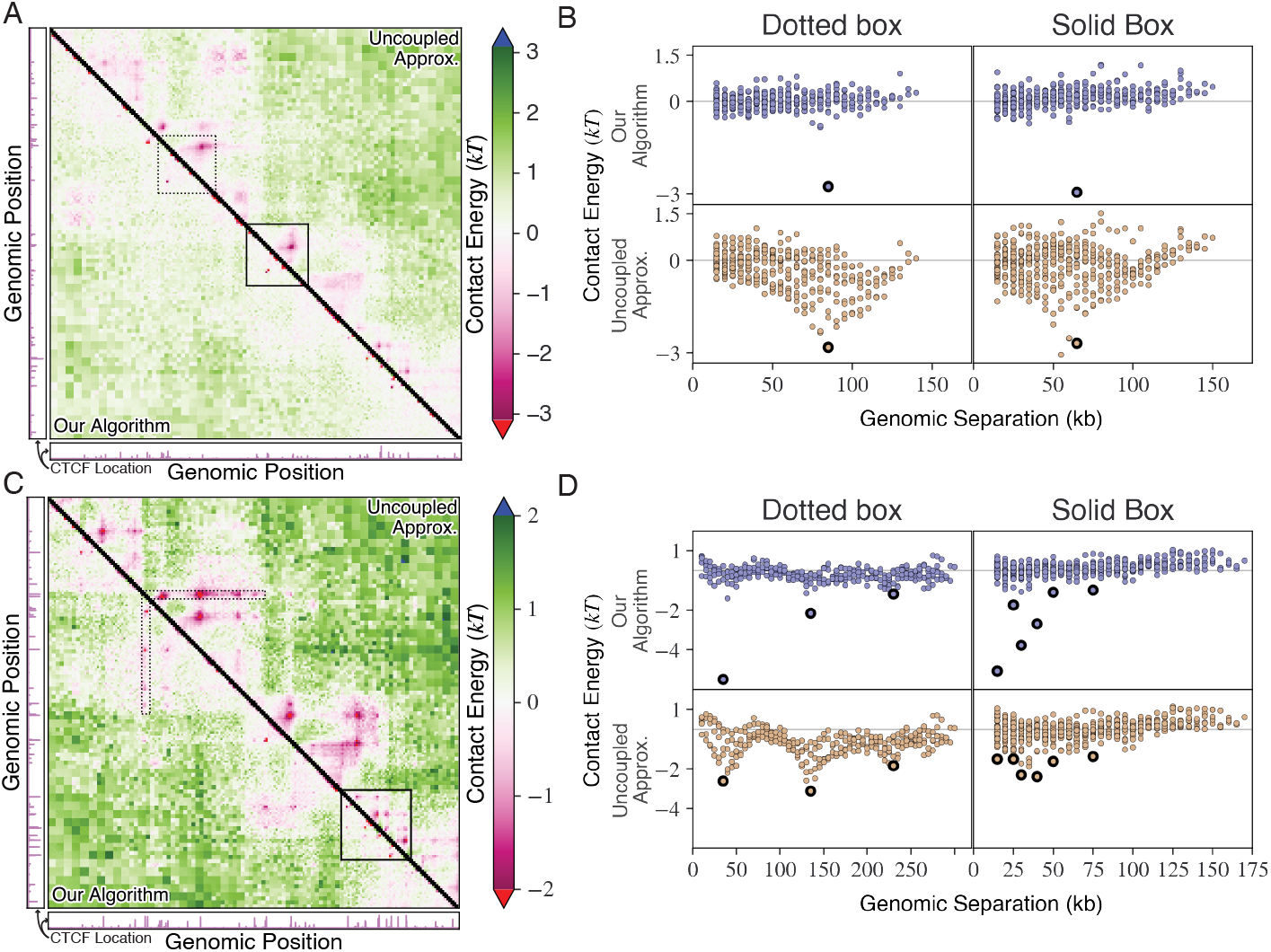
Contact energy maps contain prominent low-energy interactions, revealing the location of chromatin loop anchors. (A) Comparison of contact energies extracted with the coupled (Δ***ϵ***, lower triangle) and uncoupled (Δ***ϵ***^uc^ upper triangle) model. Each extraction used Hi-C contact probabilities from a 1 Mb region of chromosome 1, genomic positions 155, 858, 000 − 156, 858, 000, at 5 kb resolution, and the final data are displayed as such. Each boxed region contains a chromatin loop. Each axis of the contact energy maps contains a plot of the relevant CTCF ChIP-seq signals, which are independently normalized to the largest value in the plotted region. Black indicates values that are undefined in the Ising-like model. (B) Contact energies from the boxed regions in (A) are plotted against their corresponding genomic separation (distance from the map’s diagonal), with Δ***ϵ*** and Δ***ϵ***^uc^ in the upper and lower plots, respectively. Bold borders indicate the interaction with the most negative contact energy in each observed loop, as determined by our algorithm. (C) Same plot as (A) but for HFF cells. (D) Same plot as (B) but for the boxed regions in (C).

To further emphasize the difference between Δ***ϵ*** and Δ***ϵ***^uc^, Fig. 3*B* plots the energies enclosed by each box in Fig. 3*A* against the genomic separation between the relevant Hi-C bins. The contacts having the most attractive contact energies, i.e. loop anchors, are circled in black for easy identification and comparison. These presumed loop anchors have contact energies that differ only minimally from its neighbors when computed with the uncoupled approximation, whereas a significant gap can be seen between this contact energy and those associated with passive interactions when our algorithm is used. In fact, our algorithm increases the prominence of this contact from 0.280 to 0.851 and from 0.253 to 1.008 in the dotted- and solid-boxed regions, respectively. This clearly illustrates the main advantage of our algorithm, which extracts uncorrelated contact energies that facilitate the identification of mechanistically important contacts.

We note that, to highlight the contribution of each interaction to the contact probability map, we multiplied all *p*_*i*_ and 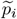 by 3 prior to computing contact energies with Eq. 6. Contact energies extracted without the multiplication factor are qualitatively similar (Fig. S5). However, increasing Hi-C contact probabilities, which are significantly smaller than the values found in homopolymer simulations, enhances the difference between the specific interaction energies for loop anchors and the non-specific ones. Scaling probabilities in this way could serve as a practical strategy to facilitate the identification of chromatin loops when analyzing contact energies. However, this additional multiplication factor is not necessary and was not used in later sections.

We analyzed the same region of HFF cells, with the corresponding results shown in Figs. 3*C* and 3*D*. Our algorithm performs well despite the overlapping loops present in this differentiated cell type, which presents a greater challenge. From left to right and compared to the uncoupled approximation, our algorithm increased the prominence values for the bold-circled contact energies from 0.384 to 0.898, from 0.347 to 0.706, and from 0.457 to 0.658 in the dotted-boxed data, while prominence increased from 0.650 to 0.947, from 0.604 to 0.924, from 0.654 to 0.999, from 0.558 to 0.910, from 0.545 to 0.894, and from 0.747 to 1.03 in the solid-boxed data. However, the magnitude of significant interactions appears to depend on loop size in these features, which we attribute to the complex interplay between nested interactions, the simplified ***J*** used to analyze their effects, and the scaling of contact probabilities.

Analyzing other regions supports the ability of contact energies to facilitate the identification of structure-driving interactions, as well (Fig. S6 and Fig. S7). Similarly, extracting contact energies from Hi-C data processed at 25 kb resolution provides largely uncorrelated results (Figs. S8 and S9), illustrating the robustness of our algorithm.

While interpreting these contact energies, it is important to recognize that the contact-space model includes first- and second-order terms only, whereas higher-order effects also influence the statistics of real polymers [48]. Of particular importance, excluded volume considerations limit the number of interactions involving each monomer. As a consequence, the probability that a monomer located in a dense region will contact a monomer located elsewhere will decrease significantly. Tracking the local density requires higher-order terms, so the Ising Hamiltonian cannot capture this effect. Therefore, to reproduce experimental contact probabilities, the Ising-like model must include a large, positive-valued correction for the free energy of interactions between monomers inside the dense region and those outside of it. This likely underlies the erroneous positive-valued contact energies associated with long-range interactions in Fig. 3*A* and Fig. 3*C*.

### Contact energies support the microphase separation of A/B compartments

At large scales, the genome is organized into self-interacting and characteristically distinct compartments [6]. Specifically, A compartments are enriched with euchromatin and active histone modifications, while B compartments are enriched with heterochromatin and silencing histone modifications [43]. These compartments may emerge from a phase separation mechanism driven by the interaction between histone modifications and potentially mediated by additional protein molecules [12, 82–107]. This mechanism requires interactions within a compartment to be more favorable than interactions between compartments, i.e. lower in energy. Contact energies extracted from Hi-C data allow a direct test of the phase separation hypothesis.

We extracted contact energies for a 100 MB region of chromosome 4 from Hi-C data of hESC and HFF cells at 500 kb resolution. In addition, we classified each interaction as AA, AB, or BB according to the compartment associated with each relevant genomic locus (see Methods section). Fig. 4*B* shows the distribution of contact energies for each interaction type, which clearly indicate that intra-compartment interactions (AA and BB) are generally favorable while inter-compartment interactions (AB) are generally unfavorable. This difference penalizes the mixing of different types of chromatin and supports a phase separation mechanism in compartment formation. Compared to hESC cells, BB interactions are more favorable, AB interactions are less favorable, and AA interactions are similarly favorable in HFF cells. Therefore, interactions involving heterochromatin appear to underlie the increased compaction of heterochromatin and stricter separation between compartment types in differentiated cells.

**Figure 4.**
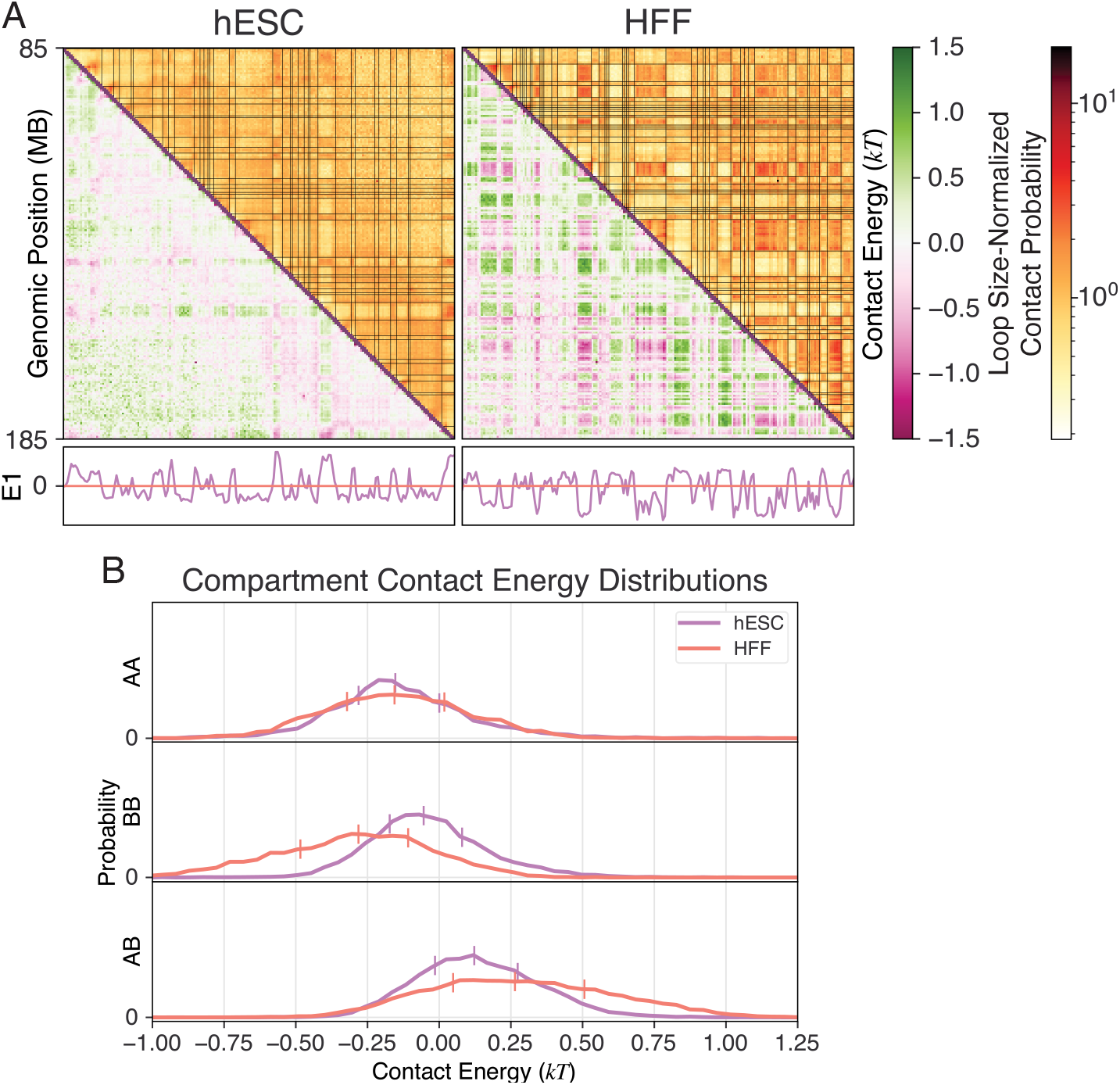
Contact energies indicate that intra-compartment interactions are more favorable than inter-compartment interactions. (A) Using the coupled model, contact energies (Δ***ϵ***, lower triangle) were extracted at 500 kb resolution in the 100 Mb region spanning genomic positions 85, 000, 000-185, 000, 000 in chromosome 4 of H1 hESC (left) and HFF (right) cells. Using eigenvector decomposition, the sign of the first eigenvector (E1) indicates compartment identity for each genomic locus, and each fine black line on the loop size-normalized Hi-C contact probability maps (upper triangle) indicates a boundary between loci in different compartments. (B) Contact energies that correspond to intra- and inter-compartment interactions maintain different probability distributions.

Compartment-level contact energies in chromosome 2 at 500 (Fig. S10) and 250 kb resolution (Fig. S11) support the same conclusion.

### Parameterizing MD simulations with extracted energies

The results presented so far suggest that extracting contact energies from Hi-C data is useful to visualize interactions and analyze structural features. At the same time, contact energies are the contact-space analogue of the interaction energies needed to parameterize accurate polymer models of chromatin, denoted *α*_*kl*_ ∈ ***α*** in Eq. 1, so the contact energies extracted by our algorithm approximate ***α***. Existing approaches [20–23] to determine ***α*** suffer from computational complexity, so parameterizing polymer models with contact energies offers a valuable alternative when using computer simulations to predict the three-dimensional structures that contribute to Hi-C probability maps. To showcase the usefulness of contact energies in this task, we parameterized polymer models with contact energies, simulated their contact probability maps, and compared the results to the initial Hi-C data.

Unlike the homopolymer model we simulated to obtain the topology-driven contact statistics, these polymer models of chromatin apply the Lennard-Jones potential defined in Eq. 15 between all monomers. As shown in Fig. 5*A*, including the excluded volume effect yields better agreement between probability decay in the *in silico* and *in vivo* contact statistics, so these models require interaction energies that account for local perturbations alone. We encoded this information in the model by using Eq. 14 with *α*_*kl*_ = 0.2Δ*ϵ*_*i*_ to apply specific interactions. Scaling contact energies by 0.2 helps correct for the approximations made by the contact-space representation of the polymer model; for example, Eq. 5 implicitly assumes that the Heaviside function in Eq. 2 defines the shape of interaction potentials, while Eq. 14 defines them with a hyperbolic tangent function. Fig. S12 analyzes the accuracy of simulated contact probability maps when contact energies are scaled by other factors.

**Figure 5.**
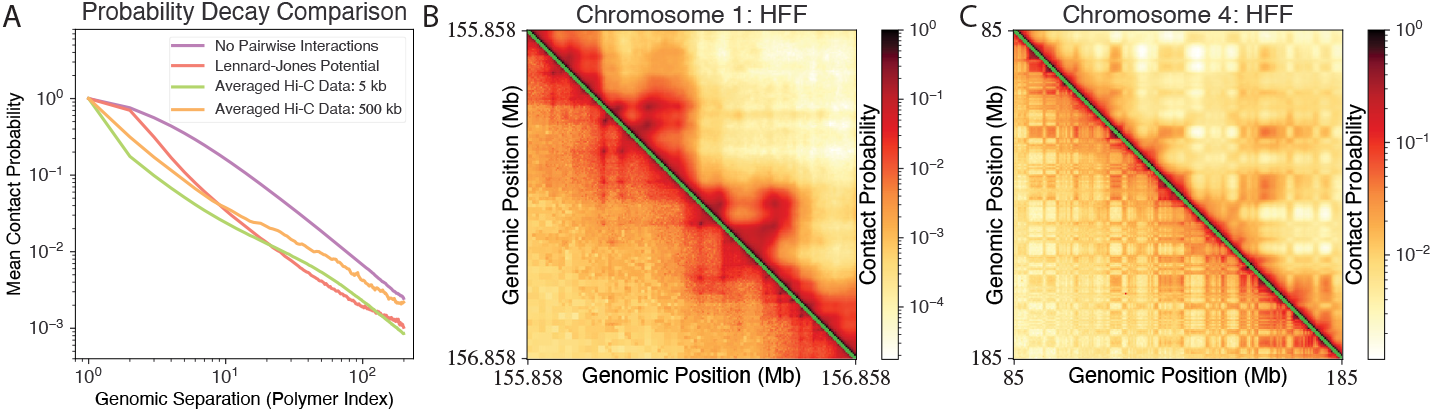
Polymer models parameterized with contact energies produce three-dimensional structures that recreate Hi-C contact maps. (A) This compares the separation dependence of the homopolymer model used to compute ***h*** and ***J***, the same model with Lennard-Jones potentials applied uniformly, and the separation-averaged Hi-C contact probabilities from chromosome 1 and 4 of HFF cells at 5 and 500 kb resolution, respectively. (B, C) These compare the experimental (lower triangle) and simulated (upper triangle) contact probabilities for HFF cells in the genomic regions (B) 155, 858, 000 − 156, 858, 000 in chromosome 1 at 5 kb resolution and (C) 85, 000, 000 − 185, 000, 000 in chromosome 4 at 500 kb resolution.

Fig. 5 presents the simulated contact probabilities for two 200-monomer models representing HFF cells at 5 (Fig. 5*B*) and 500 kb (Fig. 5*C*) resolution. These span the same regions analyzed in Fig. 3*C* and Fig. 4*A*, respectively. The first model highlights chromatin loops, while the second highlights compartments, two prominent features of chromatin organization. Visually, each model’s statistics maintain remarkable agreement with the initial Hi-C probability map, and all major features appear in the simulated data. The Pearson correlation coefficient (PCC) between the simulated and *in vivo* Hi-C contact probabilities quantifies their agreement, and the high- and low-resolution models maintain a PCC of 0.9136 and 0.8621, respectively, corroborating the visual analysis. Nevertheless, several visible artifacts contribute to the imperfect agreement between *in vivo* and *in silico* contact probabilities.

In particular, long-range contacts are less probable in the simulated models than in the *in vivo* Hi-C data. Regarding Fig. 5*C*, we primarily attribute this artifact to difference between the contact probabilities affecting the relevant homopolymer model, with Lennard-Jones interactions, and the loop size-averaged contact probabilities at 500 kb resolution (see Fig. 5*A*). On the other hand, we believe that the positive-valued contact energies associated with long-range interactions in the 5 kb-resolution model (lower triangle of Fig. S4*C*) contribute more significantly to the underprediction of long-range contact formation in Fig. 5*B*. As addressed in the discussion of Fig. 3, the contact-space model uses these positive-valued contact energies to implicitly account for higher-order topological effects that disfavor long-range interactions. Meanwhile, Cartesian-space polymer models explicitly account for this higher-order topology, so they double count its influence when directly parameterized by contact energies. As a result, simulating these polymer models underpredicts the frequency of these interactions.

These results indicate that contact energies can effectively parameterize polymer models of chromatin. Simultaneously, they confirm that the contact energies analyzed in prior subsections are physically meaningful. If more accurate predictions of chromatin structures are desired, the preexisting optimization algorithms [20–23] can use contact energies as an intelligent first guess. Given the quality of the predictions shown in Fig. 5, we anticipate that these algorithms will converge to a solution quickly when models are initialized with contact energies, thus minimizing their primary drawback.

## Conclusion

We introduced an efficient algorithm to convert correlated Hi-C contact probabilities to uncorrelated contact energies, which quantify the strength of local interactions. Our approach simplified this task by utilizing a binary contact space, where an easily invertible, Ising-like model represents polymer models of chromatin. Importantly, the Ising-like Hamiltonian separately accounts for the local interactions of interest and the system-wide effects of polymer topology. We showed that, thanks to this property, prominent contact energies correspond to loop-stabilizing interactions in *in silico* models, providing intuitive insight to the location of these structure-stabilizing interactions.

Furthermore, simulating polymer models parameterized with the extracted contact energies can predict an ensemble of three-dimensional configurations that reproduce Hi-C contact probabilities. The computational cost associated with energy extraction is nearly negligible when compared to traditional algorithms that must iteratively perform polymer simulations. This efficiency dramatically improves the accessibility of well-parameterized polymer models of chromatin, thereby democratizing the use of polymer simulations to determine, analyze, and compare the structural ensembles for chromatin regions in different cell types. In turn, these comparative structural analyses may provide valuable insight regarding the changes to chromatin organization that contribute to cell differentiation or disease progression.

Several aspects of the model can be improved. For example, as addressed in our discussion of Fig. 3, the Ising-like Hamiltonian cannot account for higher-order interactions, precluding its ability to fully capture the effects of excluded volume and other topological constraints. This limitation can cause artifacts to appear in the set of contact energies extracted in regions with multiple strong interactions, such as the positive-valued contact energies associated with long-range interactions in Fig. 3*C*. Neural network-based generative models may better approximate the probability distribution in contact space [108–110] and could improve the quality of the extracted contact energies.

Furthermore, we used Gaussian chain statistics while parameterizing the contact-space model, which likely oversimplifies the topological conditions of chromatin. Therefore, using models that better approximate chromatin systems should better account for polymer topology while extracting contact energies. Finally, the size of ***J*** is 𝒪(*M* ^4^), where *M* is the number of monomers. This presents memory challenges for larger systems, limiting the algorithm’s ability to analyze large regions at high resolution.

## Supporting information

supporting_material_final

## Author Contributions

Designed research: G.S., X.D., and B.Z.; developed new equations: G.S.; performed simulations: G.S. and B.Z.; analyzed data: G.S.; interpreted results: G.S., X.D., and B.Z.; drafted manuscript: G.S. and B.Z.; revised manuscript: G.S. and B.Z.

## Declaration of Interests

The authors declare no competing interests.

Acknowledgments

This work was supported by the National Institutes of Health (Grant R35GM133580). We thank A. Sood for many helpful conversations.

## Supporting Material

An online supplement to this article can be found by visiting BJ Online.

## Supporting Citations

References [111–113] appear in the Supporting Material.

