## supporting_material_final for "Efficient Hi-C inversion facilitates chromatin folding mechanism discovery and structure prediction"

**1 Department of Chemistry, Massachusetts Institute of Technology, Cambridge, MA 02139, USA**

*****

#### Deriving A Physically Meaningful Inversion for Strongly Coupled Models

The contact energy-extraction algorithm introduced in this work operates by inverting a contact-space representation of polymer models. Though we are most interested in polymer models of chromatin, we first considered a homopolymer model that is only affected by the topology of the system. To capture the influence of these topological constraints in contact space, we used the Ising-like Hamiltonian in Eq. 4 of the main text, i.e.

$$H_{\mathrm{hp}}(\boldsymbol{q}) = \sum_i h_i q_i + \sum_i \sum_{j>i+1} J_{ij} q_i q_j, \tag{S1}$$

to approximate the free energy associated with different homopolymer configurations. Here, $i, j \in \{1, \ldots, N\}$ index the $N = (M-1)(M-2)/2$ unique contact pairs contained in the polymer configuration $\boldsymbol{q} \equiv \{q_i\}$, where $M$ is the total number of monomers. As mentioned in the main text, we derived new inverse equations for the parameter sets $\boldsymbol{h} \equiv \{h_i\}$ and $\boldsymbol{J} \equiv \{J_{ij}\}$ to ensure that $H_{\mathrm{hp}}(\boldsymbol{q})$ faithfully captures the influence of polymer topology on contact statistics. The main text introduces the final result in Eqs. 7 and 8, which compute the model parameters using the first- and second-order homopolymer statistics $\widetilde{p}_i \equiv \langle q_i \rangle_{\mathrm{hp}}$ and $\widetilde{\mathscr{C}}_{ij} \equiv \langle q_i q_j \rangle_{\mathrm{hp}}$. In the following, we introduce a theory of total coupling in Ising-like models to explain why using the TAP inversion to extract contact energies yielded unphysical results. We then use this theory to derive Eq. 7, which determines $J_{ij}$. Afterwards, we introduce our derivation for Eq. 8, which determines $h_i$, and extend this equation to the contact-space representation of chromatin models, i.e.

$$H_{\mathrm{ME}}(\boldsymbol{q}) = \sum_i (h_i + \epsilon_i) q_i + \sum_i \sum_{j>i+1} J_{ij} q_i q_j, \tag{S2}$$

to determine the set of contact energies, $\boldsymbol{\epsilon} \equiv \{\epsilon_i\}$.

##### The TAP inversion infers unphysical contact energies

With the contact-space polymer Hamiltonian now defined, efficiently determining its parameters from contact statistics was the next necessary task while developing our algorithm. To this end, the often-used TAP inversion [1, 2] provides an efficient, closed-formed inversion of generalized Ising models in contact space. When extended to the contact energy-extraction task, this inversion determines all parameters in Eqs. S1 and S2 with

$$\begin{aligned}
J_{ij}^{\mathrm{TAP}} &= \frac{2\left(\widetilde{C}^{-1}\right)_{ij}}{1+\sqrt{1-8\left(\widetilde{C}^{-1}\right)_{ij}\left(\widetilde{p}_i-\frac{1}{2}\right)\left(\widetilde{p}_j-\frac{1}{2}\right)}} \\
h_i^{\mathrm{TAP}} &= \ln\left(\frac{1-\widetilde{p}_i}{\widetilde{p}_i}\right)-\sum_{j\neq i} J_{ij}^{\mathrm{TAP}}\widetilde{p}_j-\left(\widetilde{p}_i-\tfrac{1}{2}\right)\sum_{j\neq i}\left(J_{ij}^{\mathrm{TAP}}\right)^2\widetilde{p}_j(1-\widetilde{p}_j) \\
\epsilon_i^{\mathrm{TAP}} &= \ln\left(\frac{1-p_i}{p_i}\right)-\sum_{j\neq i} J_{ij}^{\mathrm{TAP}}p_j-\left(p_i-\tfrac{1}{2}\right)\sum_{j\neq i}\left(J_{ij}^{\mathrm{TAP}}\right)^2 p_j(1-p_j)-h_i^{\mathrm{TAP}},
\end{aligned} \tag{S3}$$

where $\widetilde{C}$ is the coveriance matrix such that $\widetilde{C}_{ij}=\widetilde{\mathscr{C}}_{ij}-\widetilde{p}_i\widetilde{p}_j$ and $\boldsymbol{p}\equiv\{p_i\}$ is the set of Hi-C contact probabilities in the chromatin system of interest. However, this approach yielded unphysical contact energies.

This failure is plainly visible in Fig. S1*I*-*L*, which compares the interaction energies used to parameterize several *in silico* models of chromatin ($\boldsymbol{\alpha}\equiv\{\alpha_{kl}\}$ from Eq. 10 of the main text, upper triangle) to the contact energies extracted using the TAP inversion ($\boldsymbol{\epsilon}$, lower triangle). In each model, we applied an attraction between two monomers to promote the formation of a single loop, and different models favored loops of different sizes; we following the same simulation protocol as in the main text to generate statistics for each *in silico* model (lower triangle of Fig. S1*A*-*D*). At loop size 22 (Fig. S1*I*), the contact energy at the contact experiencing attraction is nearly correct, but the inversion predicts that positive-valued contact energies must be applied to the surrounding contacts to reproduce the simulated contact probabilities. As the loop size associated with the promoted contact increases from 22 (Fig. S1*I*) to 50 (Fig. S1*J*), 76 (Fig. S1*K*), and 100 (Fig. S1*L*), the contact energy inferred for the promoted contact becomes worse, and the errant positive values become increasingly positive. In fact, the TAP inversion predicts that the increased contact probabilities associated with the loop size-100 case correspond to a strong repulsion between many monomers, including where the loop anchor is located, which directly contradicts the increased favorability of these interactions.

We believe that Eq. S3 fails to provide physical results because the Sherrington-Kirkpatrick (SK) model [3], for which the TAP inversion is well-suited [1,2], poorly approximates our system. In particular, mean-field inversions such as the TAP inversion perform poorly in strongly coupled systems [4]. Meanwhile, larger loops are more strongly correlated with nearby interactions than smaller loops [5,6], which agrees with the larger error in contact energies extracted with Eq. S3 when analyzing larger loops. While investigating this issue, we developed a parameter that approximates the combined influence of direct and indirect coupling on the correlation between various $q_i$.

#### Correlating energies account for both direct and indirect coupling

Correctly accounting for the combined effect of direct and indirect coupling is critically important in the contact energy-extraction task, as the strong coupling between proximal contacts implies that a driving interaction associated with $q_i$ significantly alters the contribution of $J_{j\ell}$ to $\langle q_j\rangle$ whenever both $q_j$ and $q_\ell$ are strongly correlated with $q_i$ and $J_{j\ell}$ is non-negligible; this is true regardless of the mechanistic importance of $q_j$ and $q_\ell$. To better understand this effect, we considered the effective field parameter sometimes used to investigate Ising-like models in non-equilibrium conditions [2], defined as

$$\Theta_i(\boldsymbol{q}_{\setminus i})\equiv H_{\mathrm{hp}}(\boldsymbol{q}_{\setminus i},q_i=1)-H_{\mathrm{hp}}(\boldsymbol{q}_{\setminus i},q_i=0)=h_i+\sum_{j\neq i}J_{ij}q_j \tag{S4}$$

where $\boldsymbol{q}_{\setminus i}\equiv\{q_j\forall j\neq i\}$. Conceptually, $\Theta_i$ determines the total change in free energy that occurs when contact $i$ is formed from some starting configuration, and we relate it to homopolymer contact probabilities with

$$\begin{aligned}
\widetilde{p}_i &= \widetilde{p}_j\left\langle q_i\right\rangle_{q_j=1}+(1-\widetilde{p}_j)\left\langle q_i\right\rangle_{q_j=0} \\
&= \widetilde{p}_j\left\langle\frac{1}{1+\exp\left(\Theta_i(\boldsymbol{q}_{\setminus i})\right)}\right\rangle_{q_j=1}+(1-\widetilde{p}_j)\left\langle\frac{1}{1+\exp\left(\Theta_i(\boldsymbol{q}_{\setminus i})\right)}\right\rangle_{q_j=0}.
\end{aligned} \tag{S5}$$

Here, $\langle X\rangle_Y$ denotes a conditional expectation, so $\langle q_i\rangle_{q_j=1} = \widetilde{\mathscr{C}}_{ij}/\widetilde{p}_j$ and $\langle q_i\rangle_{q_j=0} = (\widetilde{p}_i - \widetilde{\mathscr{C}}_{ij})/(1-\widetilde{p}_j)$. Therefore, using all $j \neq i$, $(N-1)$ analogs of Eq. S5 relate $h_i$ and $J_{ij}\forall j \neq i$ to the contact statistics they must capture ($\widetilde{p}_i$ and $\widetilde{\mathscr{C}}_{ij}\forall j \neq i$).

Similarly, the difference $U_\mathrm{c}(i,j) \equiv \left\langle \Theta_i(\boldsymbol{q}_{\setminus i})\right\rangle_{q_j=1} - \left\langle \Theta_i(\boldsymbol{q}_{\setminus i})\right\rangle_{q_j=0}$ quantifies the effect contact $j$'s existence has on contact $i$'s favorability, so we refer to it as the *correlating energy* between $q_i$ and $q_j$. $h_i$ cancels in $U_\mathrm{c}(i,j)$, which relates $J_{ij}\forall j \neq i$ to the relevant second-order statistics:

$$U_\mathrm{c}(i,j) = J_{ij} + \sum_{\ell \neq i,j} J_{i\ell}\frac{\widetilde{C}_{j\ell}}{\widetilde{C}_{jj}} \approx \ln\left(\frac{\left(\widetilde{p}_i - \widetilde{\mathscr{C}}_{ij}\right)\left(\widetilde{p}_j - \widetilde{\mathscr{C}}_{ij}\right)}{\widetilde{\mathscr{C}}_{ij}\left(1-\widetilde{p}_i-\widetilde{p}_j+\widetilde{\mathscr{C}}_{ij}\right)}\right) \equiv U'_\mathrm{c}(i,j). \tag{S6}$$

We derived the right-hand side of Eq. S6 by applying the mean-field approximation, i.e. $\langle f(x)\rangle \approx f(\langle x\rangle)$, to the conditional expectations in Eq. S5 and solving for $\left\langle \Theta_i(\boldsymbol{q}_{\setminus i})\right\rangle_{q_j=0/1}$ with respect to conditional probabilities.

For each model investigated in Fig. S1, Fig. S1*E*-*H* compare the correlating energies approximated with statistics ($U'_\mathrm{c}(i,j)$, upper triangle) to the correlating energies computed with the TAP model ($U_\mathrm{c}(i,j)$, lower triangle). While the correlating energy computed with statistics matches correlations in the model (Fig. S1*A*-*D*, upper triangle), the TAP model predicts that the formation of loops – especially larger loops – *increases* the energy associated with many contacts. For each loop size, the artifacts in the correlating energies strongly agree with the artifacts in the contact energies (Fig. S1*I*-*L*), corroborating their relevance.

#### Statistics-based correlating energies are physical

Though correlating energies appear to be related to the success or failure of contact energy extraction when using the TAP inversion, we wanted to confirm that the right-hand side of Eq. S6 is physically accurate. To do this, we used pseudolikelihood maximization (PLM), discovered independently in 1974 [7] and 2012 [4], to determine $\boldsymbol{J}^\mathrm{PLM}$. As the name suggests, this approach maximizes an approximation of the likelihood function, so it computes model parameters that approximately maximize entropy. This approximation works well in the case of strong coupling [4], so PLM can determine physically accurate coupling parameters for the homopolymer model in contact space. Knowing this, we used PLM to compute $\boldsymbol{J}^\mathrm{PLM}$, then we validated the right-hand side of Eq. S6 by comparing its predictions to correlating energies computed using $\boldsymbol{J}^\mathrm{PLM}$, $U_\mathrm{c}^\mathrm{PLM}(i,j)$.

PLM avoids the (log-)likelihood function, which is computationally intractable [4, 7, 8], by approximating the log-likelihood function with [8]

$$L_{\boldsymbol{Q}}^{(0)}\left(\boldsymbol{h}^\mathrm{PLM}, \boldsymbol{J}^\mathrm{PLM}\right) = \sum_{\boldsymbol{q}\in\boldsymbol{Q}}\sum_{i=1}^{N}\log\left[\mathbb{P}\left(q_i|\boldsymbol{q}_{\setminus i}, \boldsymbol{h}, \boldsymbol{J}\right)\right] = \sum_{\boldsymbol{q}\in\boldsymbol{Q}}\sum_{i=1}^{N}\left[1+\exp\left\{(2q_i-1)\left(h_i^\mathrm{PLM} + \sum_{j\neq i} J_{ij}^\mathrm{PLM} q_j\right)\right\}\right]^{-1}, \tag{S7}$$

where $\boldsymbol{Q}$ is the known set of homopolymer configurations. (Due to memory constraints, $\boldsymbol{Q}$ consists of 50,000 from the total 200,000 homopolymer configurations computed in the main text.) Parameters inferred by PLM suffer from overfitting unless they are regularized [8], so we added the square of the $l_2$ norm for each $J_{ij}^\mathrm{PLM}$ to Eq. S7, as suggested in a prior application of PLM [8]. Thus, rather than directly maximizing Eq. S7, we minimized

$$-L_{\boldsymbol{Q}}\left(\boldsymbol{h}^\mathrm{PLM}, \boldsymbol{J}^\mathrm{PLM}\right) = -L_{\boldsymbol{Q}}^{(0)}\left(\boldsymbol{h}^\mathrm{PLM}, \boldsymbol{J}^\mathrm{PLM}\right) + \lambda_J\sum_i\sum_j\left(J_{ij}^\mathrm{PLM}\right)^2 \tag{S8}$$

and scaled the regularization term by $\lambda_J = 0.2$ to avoid unreasonably small $J_{ij}^\mathrm{PLM}$ values while still preventing overfitting. We performed this computation in Julia 1.5.2 [9] using a publicly available implementation of PLM [10].

Fig. S2 compares the correlating energies computed with $\boldsymbol{J}^\mathrm{PLM}$ ($U_\mathrm{c}^\mathrm{PLM}(i,j)$, lower triangle) to those approximated with statistics only ($U'_\mathrm{c}(i,j)$, upper triangle). Both calculations qualitatively agree regardless of the loop size of the analyzed contact. The primary difference is the magnitude of each correlating energy: $\left|U_\mathrm{c}^\mathrm{PLM}(i,j)\right|$ slightly decreases in magnitude as loop size increases, while $|U'_\mathrm{c}(i,j)|$ increases with loop size.

However, larger loops are associated with stronger correlations (upper triangle of Fig. S1*A*-*D*) [5, 6], so their correlating energies should be more negative. This artifact in $U_{\mathrm{c}}^{\mathrm{PLM}}(i,j)$ results from the regularization term in Eq. S8, which favors small $\left|J_{ij}^{\mathrm{PLM}}\right|$, and the limited size of $\boldsymbol{Q}$, which minimizes the contribution of the correlation between infrequent contacts (i.e. large loops) to $L_{\boldsymbol{Q}}^{(0)}$; the latter point is supported by the substantial noise present in the $U_{\mathrm{c}}^{\mathrm{PLM}}(i,j)$ associated with interactions between large loops. Therefore, while the set of $U_{\mathrm{c}}^{\mathrm{PLM}}(i,j)$ confirms that $U_{\mathrm{c}}'(i,j)$ are qualitatively correct at each contact site, we believe that $U_{\mathrm{c}}'(i,j)$ more accurately predicts the scale of correlating energies.

#### Computing coupling parameters that reproduce correlating energies

Given the physical quality of $U_{\mathrm{c}}'(i,j)$, Eq. S6 provides a convincing relationship between the known first- and second-order contact statistics and the desired $J_{ij}$ values. Conveniently, this equation can be solved to find the $\boldsymbol{J}$ that exactly reproduces the correlating energies approximated with statistics, i.e. $U_{\mathrm{c}}'(i,j)$. This yields Eq. 7 from the main text, rewritten here for convenience:

$$J_{ij} = \sum_{\ell\neq i} \ln\left(\frac{\left(\widetilde{p}_i - \widetilde{\mathscr{C}}_{i\ell}\right)\left(\widetilde{p}_\ell - \widetilde{\mathscr{C}}_{i\ell}\right)}{\widetilde{\mathscr{C}}_{i\ell}\left(1-\widetilde{p}_i-\widetilde{p}_\ell+\widetilde{\mathscr{C}}_{i\ell}\right)}\right)\widetilde{p}_\ell\left(1-\widetilde{p}_\ell\right)\left(\left(\widetilde{C}_{\overline{ii}}\right)^{-1}\right)_{j\ell} \approx \sum_{\ell\neq i} U_{\mathrm{c}}(i,\ell)\widetilde{p}_\ell\left(1-\widetilde{p}_\ell\right)\left(\left(\widetilde{C}_{\overline{ii}}\right)^{-1}\right)_{j\ell} = J_{ij}^{\mathrm{exact}}. \tag{S9}$$

It is worth noting that computing $\boldsymbol{J}$ with Eq. S9 requires inverting $N$ submatrices of $C$, each of size $(N-1)\times(N-1)$ (computational complexity $\mathcal{O}(N^4)$), while other statistics-based approaches [1, 2] simply invert the covariance matrix once (computational complexity $\mathcal{O}(N^3)$). A math identity [11] brings the computational complexity of our approach to parity with the existing approaches, yielding

$$J_{ij} = \sum_{\ell\neq i} \ln\left(\frac{\left(\widetilde{p}_i - \widetilde{\mathscr{C}}_{i\ell}\right)\left(\widetilde{p}_\ell - \widetilde{\mathscr{C}}_{i\ell}\right)}{\widetilde{\mathscr{C}}_{i\ell}\left(1-\widetilde{p}_i-\widetilde{p}_\ell+\widetilde{\mathscr{C}}_{i\ell}\right)}\right)\widetilde{p}_\ell\left(1-\widetilde{p}_\ell\right)\left[\left(\widetilde{C}^{-1}\right)_{j\ell} - \frac{\left(\widetilde{C}^{-1}\right)_{ji}\left(\widetilde{C}^{-1}\right)_{i\ell}}{\left(\widetilde{C}^{-1}\right)_{ii}}\right]. \tag{S10}$$

Nonetheless, we used Eq. S9 to compute $\boldsymbol{J}$ in this work because implementing Eq. S10 presented numerical challenges related to the precision of $\widetilde{C}^{-1}$. In the future, using high-precision variables to invert $\widetilde{C}$ will allow our inversion to be implemented in the form of Eq. S10, greatly reducing the computational complexity associated with Eq. S9 and allowing much larger systems to be analyzed.

#### Computing first-order model parameters with contact statistics

As with $\boldsymbol{J}$, we developed an inverse equation to compute the first-order parameters $h_i \in \boldsymbol{h}$ and $\epsilon_i \in \boldsymbol{\epsilon}$. The derivation starts by manipulating the equality $\left\langle \exp\left(-\Theta_i(\boldsymbol{q}_{\setminus i})\right)\right\rangle_{q_i=0} = \widetilde{p}_i/(1-\widetilde{p}_i)$ to find

$$h_i = \ln\left(\frac{1-\widetilde{p}_i}{\widetilde{p}_i}\right) + \ln\left\langle \exp\left(-\sum_{j\neq i} J_{ij}q_j\right)\right\rangle_{q_i=0}. \tag{S11}$$

Manipulating $\left\langle \exp\left(+\Theta_i(\boldsymbol{q}_{\setminus i})\right)\right\rangle_{q_i=1} = (1-\widetilde{p}_i)/\widetilde{p}_i$ yields a similar result. We combine these representations of $h_i$ to find

$$h_i = \ln\left(\frac{1-\widetilde{p}_i}{\widetilde{p}_i}\right) + (1-\widetilde{p}_i)\ln\left\langle \exp\left(-\sum_{j\neq i} J_{ij}q_j\right)\right\rangle_{q_i=0} - \widetilde{p}_i\ln\left\langle \exp\left(+\sum_{j\neq i} J_{ij}q_j\right)\right\rangle_{q_i=1}, \tag{S12}$$

where Eq. S11 and its analogue are weighted by the probability of the condition each imposes. Next, we approximate $\mathbb{P}\left(\boldsymbol{q}_{\setminus i}|q_i=0\right) \approx \prod_{j\neq i}\mathbb{P}\left(q_j|q_i=0\right)$ and $\mathbb{P}\left(\boldsymbol{q}_{\setminus i}|q_i=1\right) \approx \prod_{j\neq i}\mathbb{P}\left(q_j|q_i=1\right)$ to compute the expectations in Eq. S12, yielding

$$\begin{aligned} h_i \approx \ln\left(\frac{1-\widetilde{p}_i}{\widetilde{p}_i}\right) + (1-\widetilde{p}_i)\sum_{j\neq i}\ln\left[\left(\frac{1-\widetilde{p}_i-\widetilde{p}_j+\widetilde{\mathscr{C}}_{ij}}{1-\widetilde{p}_i}\right)+\left(\frac{\widetilde{p}_j-\widetilde{\mathscr{C}}_{ij}}{1-\widetilde{p}_i}\right)e^{-J_{ij}}\right] \\ -\widetilde{p}_i\sum_{j\neq i}\ln\left[\left(\frac{\widetilde{p}_i-\widetilde{\mathscr{C}}_{ij}}{\widetilde{p}_i}\right)+\left(\frac{\widetilde{\mathscr{C}}_{ij}}{\widetilde{p}_i}\right)e^{+J_{ij}}\right]. \end{aligned} \tag{S13}$$

Eq. S13 cannot be applied to Hi-C data because Hi-C experiments do not provide second-order contact statistics. Therefore, $\mathbb{P}\left(q_j = 1 | q_i = 1\right) = \widetilde{\mathscr{C}}_{ij}/\widetilde{p}_i$ and $\mathbb{P}\left(q_j = 1 | q_i = 0\right) = (\widetilde{p}_j - \widetilde{\mathscr{C}}_{ij})/(1-\widetilde{p}_i)$ must be approximated by $\widetilde{p}_j$ and $(1-\widetilde{p}_j)$, respectively. This changes Eq. S13 into Eq. 8 from the main text, i.e.

$$h_i = \ln\left(\frac{1-\widetilde{p}_i}{\widetilde{p}_i}\right) + \sum_{j\neq i}\ln\left[\frac{\left((1-\widetilde{p}_j)+\widetilde{p}_j e^{-J_{ij}}\right)^{1-\widetilde{p}_i}}{\left((1-\widetilde{p}_j)+\widetilde{p}_j e^{+J_{ij}}\right)^{\widetilde{p}_i}}\right]. \tag{S14}$$

Eq. S14 computes $(h_i + \epsilon_i)$ when used to invert $H_{\mathrm{ME}}$, and subtracting $h_i$ from each side yields Eq. 6 in the main text, defined as

$$\epsilon_i = \ln\left(\frac{1-p_i}{p_i}\right) + \sum_{j\neq i}\ln\left[\frac{\left((1-p_j)+p_j e^{+J_{ij}}\right)^{p_i}}{\left((1-p_j)+p_j e^{-J_{ij}}\right)^{1-p_i}}\right] - h_i. \tag{S15}$$

### Minimizing Noise in the Simulated Homopolymer Contact Statistics

As mentioned in the Methods section of the main text, we minimized uncertainty in the simulated homopolymer statistics by averaging the statistics associated with topologically identical interactions. Chan and Dill's analysis of infinite homopolymers [5, 6] provided a simple framework that enabled this approach.

They used two parameters to describe the topology of interactions in an infinite homopolymer. First, indexing monomers with $k < l$, $K \equiv l - k$ describes the *loop size* associated with each contact, so contacts $i$ and $j$ with $K^{(i)} = K^{(j)}$ have the same contact probability. The correlation between distinct contacts depends on both of their loop sizes, and these contacts are indexed as 1 and 2 such that $l_1 - k_1 = K_1 \leq K_2 = l_2 - k_2$. Correlation also depends on the *separation* between these loops, defined as $L \equiv l_2 - l_1$, which affects the degree of overlap between the two loops. Accounting for reflection symmetry, the separations $L$ and $K_1 + K_2 - L$ are equivalent. As a consequence, two pairs of contacts, labeled $A \equiv (i_1, j_1)$ and $B \equiv (i_2, j_2)$, have identical second-order statistics if $K_1^{(A)} = K_1^{(B)}$, $K_2^{(A)} = K_2^{(B)}$, and either $L^{(A)} = L^{(B)}$ or $L^{(A)} = K_2^{(B)} + K_1^{(B)} - L^{(B)}$. (Other conditions imply that additional second-order statistics should be identical [5, 6], but we ignored this deeper analysis when refining the simulated contact statistics.)

With this knowledge, we collected the first- and second-order statistics for the central 300-monomer region of the homopolymer and replaced those predicted to be identical with their average. Recalling that the simulated homopolymer is 500 monomers long, this minimizes edge effects while also providing a greater reduction in uncertainty than analyzing just the central 200-monomer region of interest would. The latter point is particularly important for statistics associated with large loops. For example, the largest loop in the 200-monomer system has size $K = 199$, and the central 300-monomer region contains 101 copies of each statistic associated with it and relevant to the 200-monomer system, while the central 200-monomer region contains only 1 copy of these statistics.

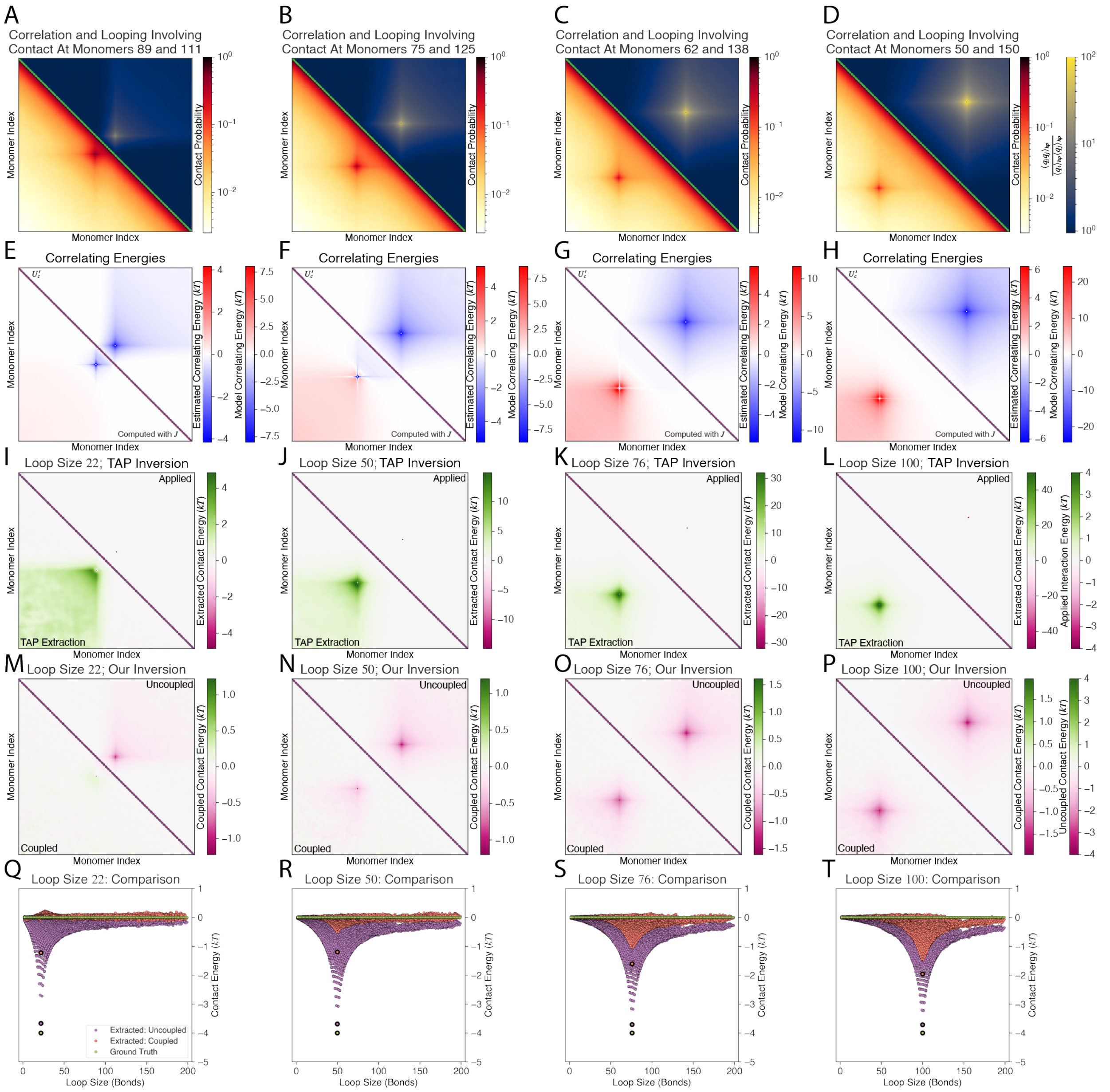


**Figure S1. Correlating energies represent the total coupling between contacts and underlie the artifacts observed in TAP-derived contact energies.** In four independent models, we applied an attraction between the two monomers involved with a single contact, thus promoting loop formation for a loop of size 22 (A,E,I,M,Q), 50 (B,F,J,N,R), 76 (C,G,K,O,S), or 100 (D,H,L,P,T). Each attraction took the form of Eq. 10 and had strength $\alpha_{kl} = -4kT$. (A-D) The simulated contact probabilities of each model (lower triangle) agree strongly with the correlation between that contact and all others (upper triangle), which can be quantified as $\widetilde{\mathscr{C}}_{ij}/(\widetilde{p}_i\widetilde{p}_j)$ [5,6]. (E-H) This result agrees well with $U_c'(i,j)$ (upper triangle), which approximates the physically ideal correlating energies. In comparison, the correlating energies produced by the TAP-derived model (lower triangle) disagree in magnitude at loop size 22 (E) and both magnitude and sign for loop sizes 50 (F), 76 (G), and 100 (H). (I-L) Compared to the $\{\alpha_{kl}\}$ parameterizing each *in silico* model (upper triangle), contact energies extracted from the model's simulated contact statistics using the TAP inversion (lower triangle) contain artifacts that are characteristically similar to those observed in TAP-derived correlating energies at loop size 22 (I), 50 (J), 76 (K), and 100 (L). (M-T) The contact energies extracted by our approach (lower triangle in M-T, orange points in Q-T) display less correlation than those extracted with an uncoupled model (upper triangle in M-P, purple in Q-T). The interaction energies applied in the polymer models are shown for comparison in the 1D plots (green points, Q-T).

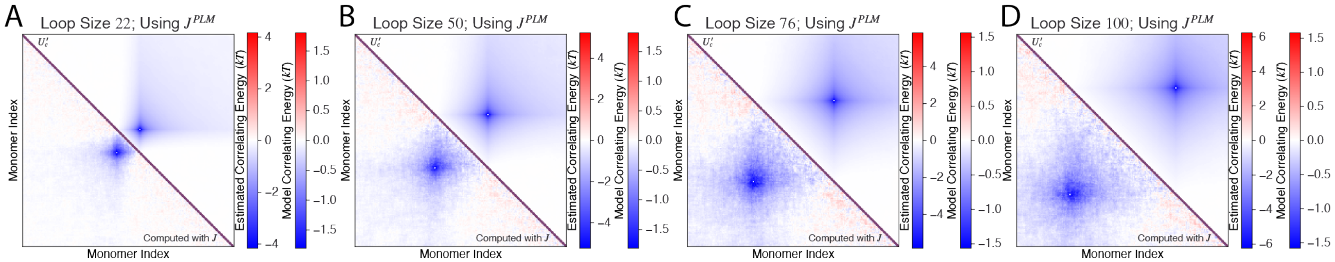


**Figure S2. Computing correlating energies with a maximum pseudolikelihood-derived model supports the physical accuracy of the correlating energies approximated using statistics.** Fixing $i$ as the index corresponding to the contact involving monomer pair $(10, 32)$ (A), $(75, 125)$ (B), $(62, 138)$ (C), and $(50, 150)$ (D), the correlating energies between contact $i$ and all other contacts are plotted in two dimensions. Characteristically, the correlating energies computed with the maximum pseudolikelihood-derived model parameters $J_{ij}^{\mathrm{PLM}} \in \boldsymbol{J}^{\mathrm{PLM}}$ (lower triangle) agree with those using homopolymer statistics to approximate the physically ideal result ($U_c'(i, j)$, upper triangle), which our model exactly reproduces. Purple indicates values that are undefined in the contact-space model.

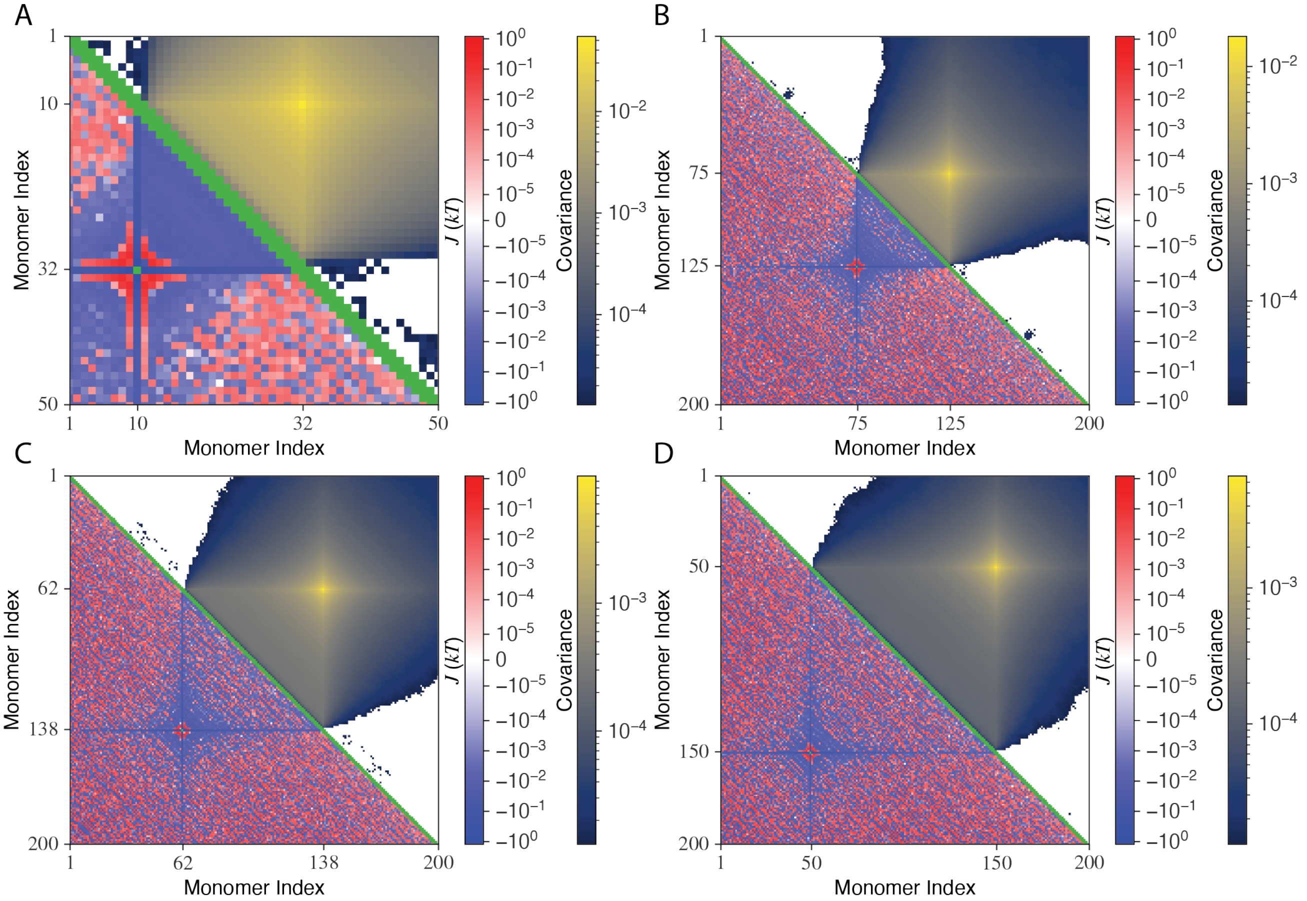


**Figure S3. The direct coupling in our model supports the correlation between contacts.** Fixing $i$ as the index corresponding to the contact involving monomer pair $(10, 32)$ (A), $(75, 125)$ (B), $(62, 138)$ (C), and $(50, 150)$ (D), the coupling parameters $J_{ij} \forall j \neq i$ (lower triangle) and covariance $\widetilde{\mathscr{C}}_{ij} - \widetilde{p}_i \widetilde{p}_j \forall j \neq i$ (upper triangle) are shown, where the latter is a measure of the correlation between contacts. Green indicates values that are undefined in the contact-space model.

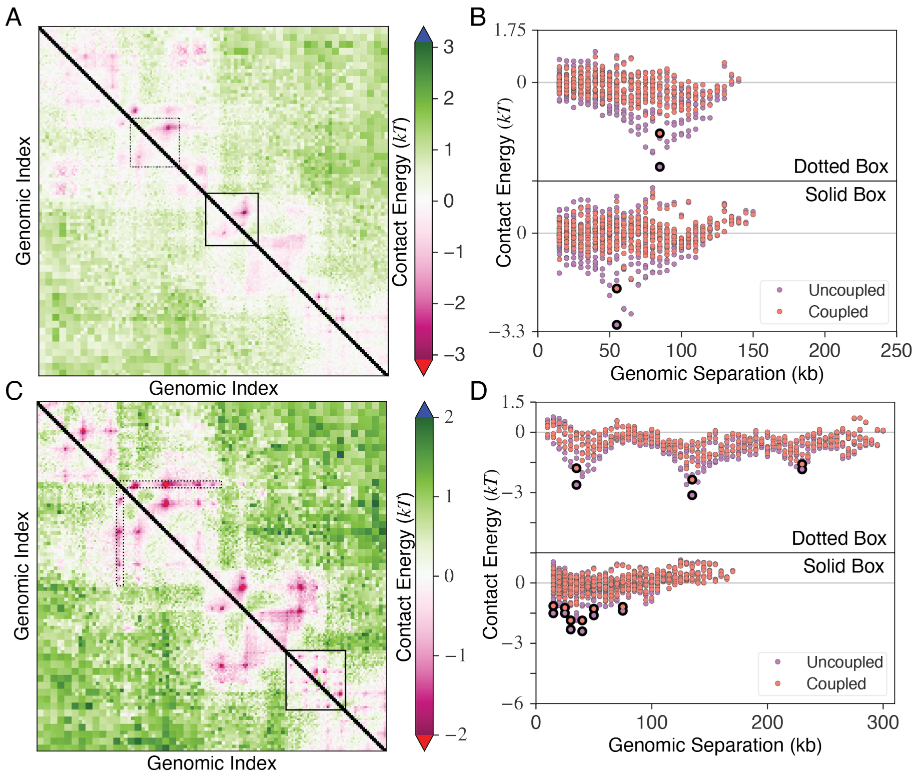


**Figure S4. Contact energies extracted from unscaled Hi-C contact probabilities maintains greater similarity with those extracted using the uncoupled model.** (A) Contact energies extracted with the coupled ($\Delta\epsilon$, lower triangle) and uncoupled ($\Delta\epsilon^{uc}$, upper triangle) models are compared. Each extraction used Hi-C contact probabilities from the 1 Mb region in chromosome 1 of H1 hESC cells, genomic position 155,858,000-156,858,000, at 5 kb resolution, and the final data are displayed as such. Each boxed region contains a chromatin loop. Black indicates values that are undefined in the contact-space model. (B) The contact energies from each boxed region in (A) are plotted against their corresponding genomic separation (distance from the map's diagonal), with $\Delta\epsilon$ in orange and $\Delta\epsilon^{uc}$ in purple. Circled points indicate the contact energies that are visibly prominent in the 2D plot (lower triangle, A). (C) Same plot as (A) but for HFF cells. (D) Same plot as (B) but for the boxed regions in (C).

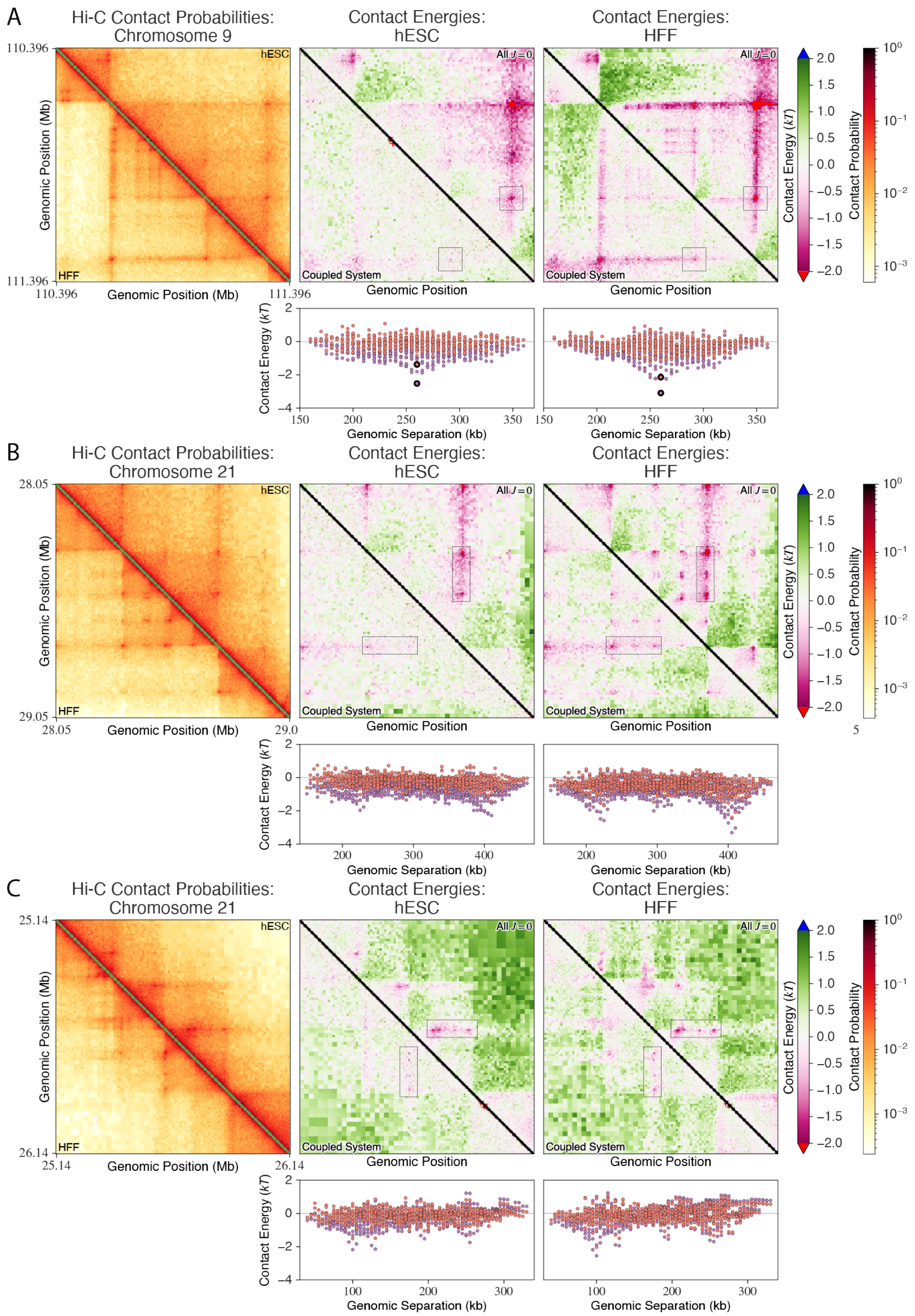


**Figure S5. Extracting contact energies from scaled Hi-C probabilities facilitates the identification of important contacts at 5 kb resolution.** (A-C) The left panel displays the Hi-C contact probabilities for the 1 Mb region spanning genomic positions 110,396,000-111,396,000 in chromosome 9 (A), 28,050,000-29,050,000 in chromosome 21 (B), or 25,140,000-26,140,000 in chromosome 21 (C), which we preprocessed according to the procedure described in the main text using the same hESC (upper triangle) and HFF (lower triangle) datasets [12]. The contact energy maps display the contact energies extracted from the same hESC (center panel) and HFF (right panel) Hi-C contact probabilities using the coupled ($\Delta\epsilon$, lower triangle) and uncoupled ($\Delta\epsilon^{\mathrm{uc}}$, upper triangle) models. A boxed region contains the contact energies near chromatin loop anchors, which are plotted against their genomic separation in the 1D plot below. We scaled the displayed and loop size-averaged Hi-C contact probabilities by 3 prior to extracting energies in the coupled case, but they remained unscaled during the uncoupled extraction. For 2D plots of contact probabilities and energies, respectively, green and black indicate values that are undefined in the contact-space model.

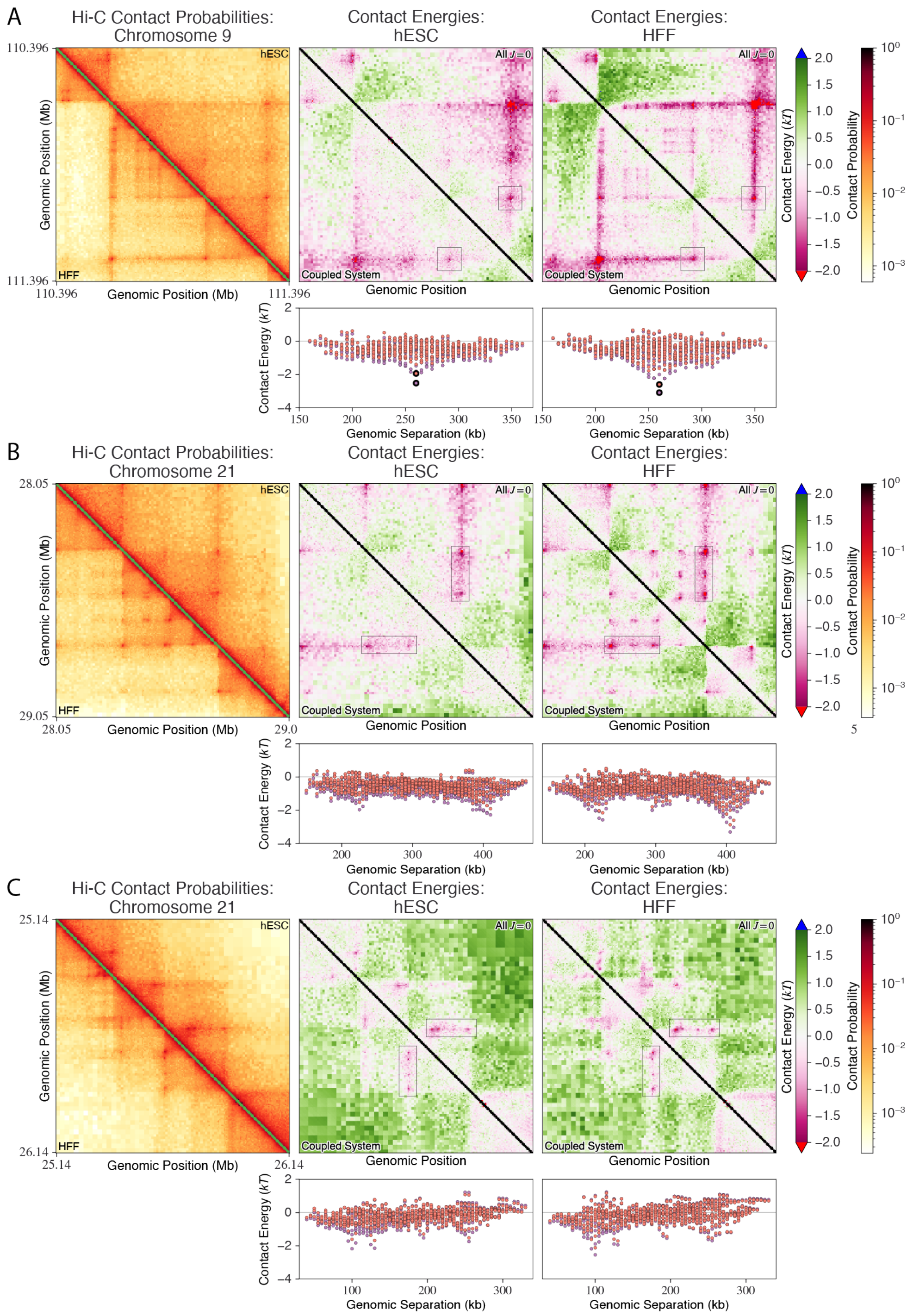


**Figure S6. Extracting contact energies from unscaled Hi-C probabilities facilitates the identification of important contacts at 5 kb resolution.** This figure is identical to Fig. S5, except all contact energies were extracted from unscaled Hi-C contact probabilities.

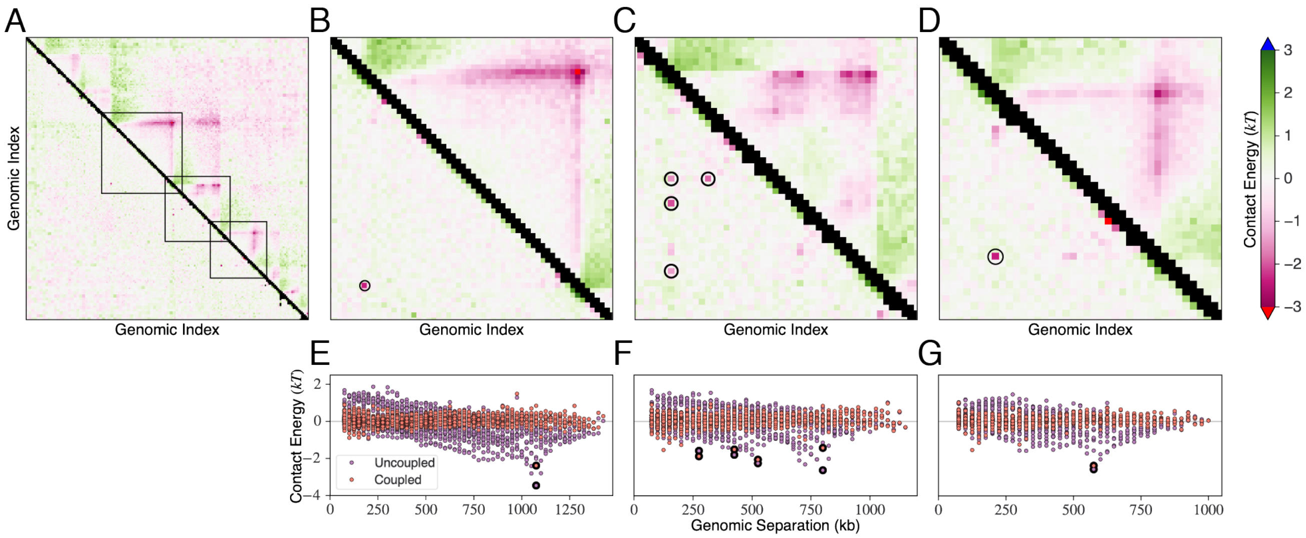


**Figure S7. Extracting contact energies with the coupled model facilitates the identification of important contacts at 25 kb resolution.** Contact energies were extracted for the 5 Mb region spanning 82440000 − 87440000 in chromosome 7 at 25 kb resolution in H1 hESC cells, and several boxed regions contain chromatin loops (A). The top left (B/E), center (C/F), and bottom right (E/G) boxed regions are highlighted separately. Two-dimensional maps display the features in the coupled ($\Delta\boldsymbol{\epsilon}$, lower triangle) and uncoupled ($\Delta\boldsymbol{\epsilon}^{\mathrm{uc}}$, upper triangle) contact energies (E-G), while one-dimensional plots emphasize the prominence of presumed structurally important interaction sites (bold circled) identified in the 2D maps (circled in lower triangle). We scaled the Hi-C contact probabilities by 3 prior to extracting energies in the coupled case, but they remained unscaled during the uncoupled extraction.

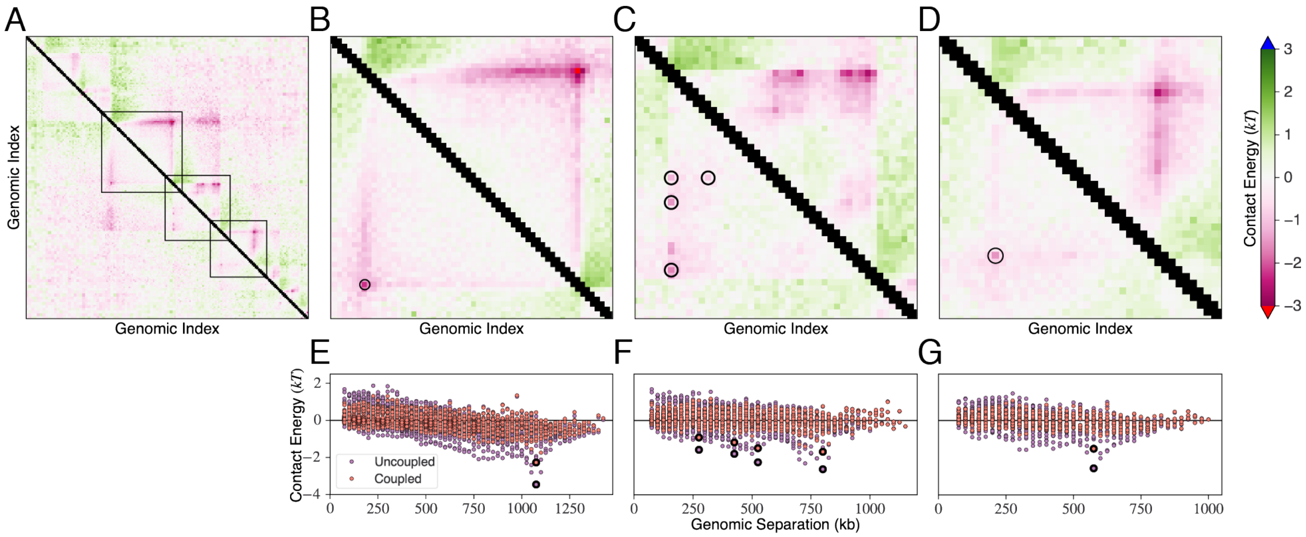


**Figure S8. Extracting contact energies with the coupled model facilitates the identification of important contacts at 25 kb resolution.** This figure is identical to Fig. S7, except all contact energies were extracted from unscaled Hi-C contact probabilities.

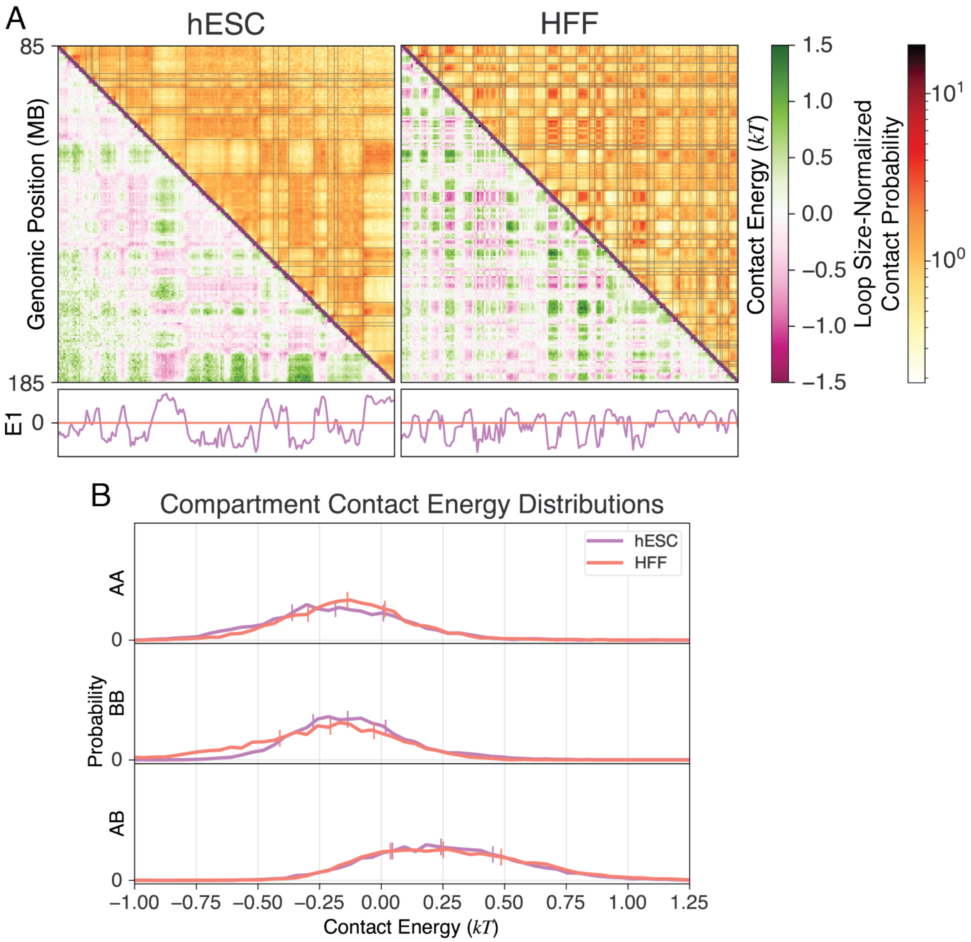


**Figure S9. Contact energies indicate that intra-compartment interactions are more favorable than inter-compartment interactions in chromosome 2.** (A) Using the coupled model, contact energies ($\Delta\epsilon$, lower triangle) were extracted at 500 kb resolution in the 100 Mb region spanning genomic positions 140,000,000-240,000,000 in chromosome 2 of H1 hESC (left) and HFF (right) cells. After performing eigenvector decomposition, the sign of the first eigenvector (E1) indicates compartment identity for each genomic locus, and each fine black line on the loop size-normalized Hi-C contact probability maps (upper triangle) indicates a boundary between loci in different compartments. (B) Contact energies that correspond to intra- and inter-compartment interactions maintain different probability distributions.

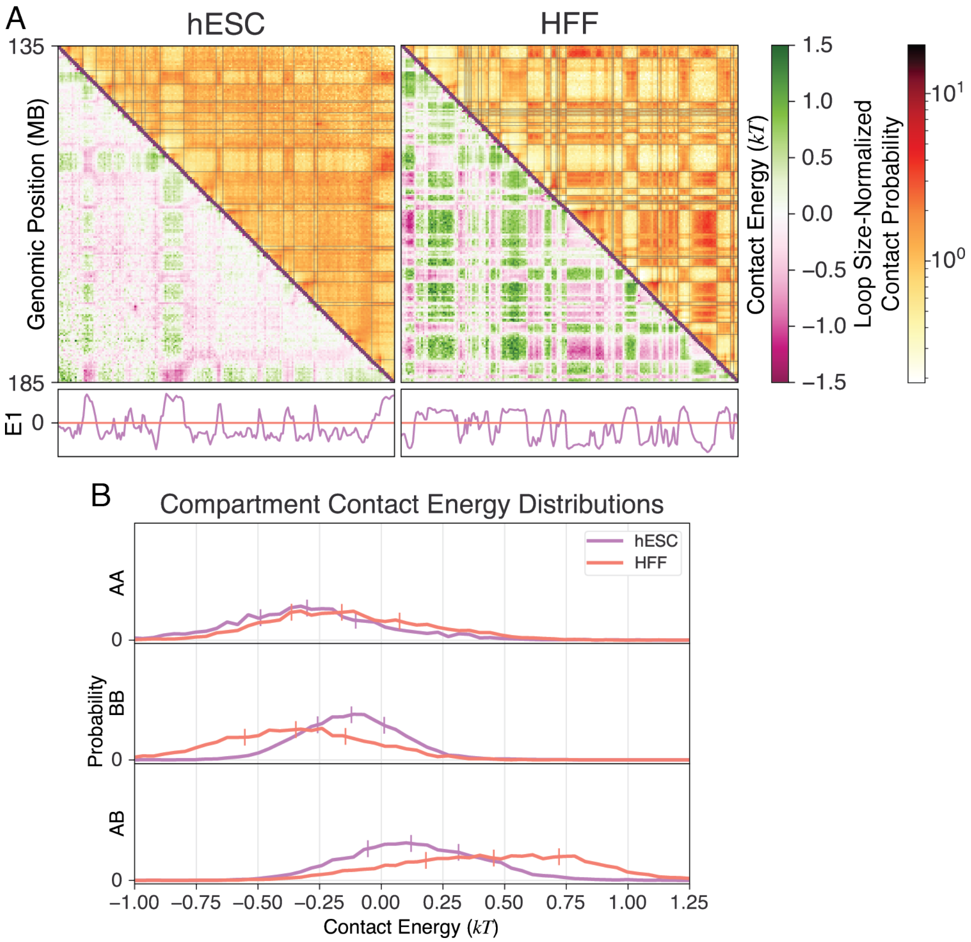


**Figure S10. Contact energies extracted at 250 kb resolution indicate that intra-compartment interactions are more favorable than inter-compartment interactions.** (A) Using the coupled model, contact energies ($\Delta\epsilon$, lower triangle) were extracted at 250 kb resolution in the 50 Mb region spanning genomic positions 135,000,000-185,000,000 in chromosome 4 of H1 hESC (left) and HFF (right) cells. After performing eigenvector decomposition, the sign of the first eigenvector (E1) indicates compartment identity for each genomic locus, and each fine black line on the loop size-normalized Hi-C contact probability maps (upper triangle) indicates a boundary between loci in different compartments. (B) Contact energies that correspond to intra- and inter-compartment interactions maintain different probability distributions.

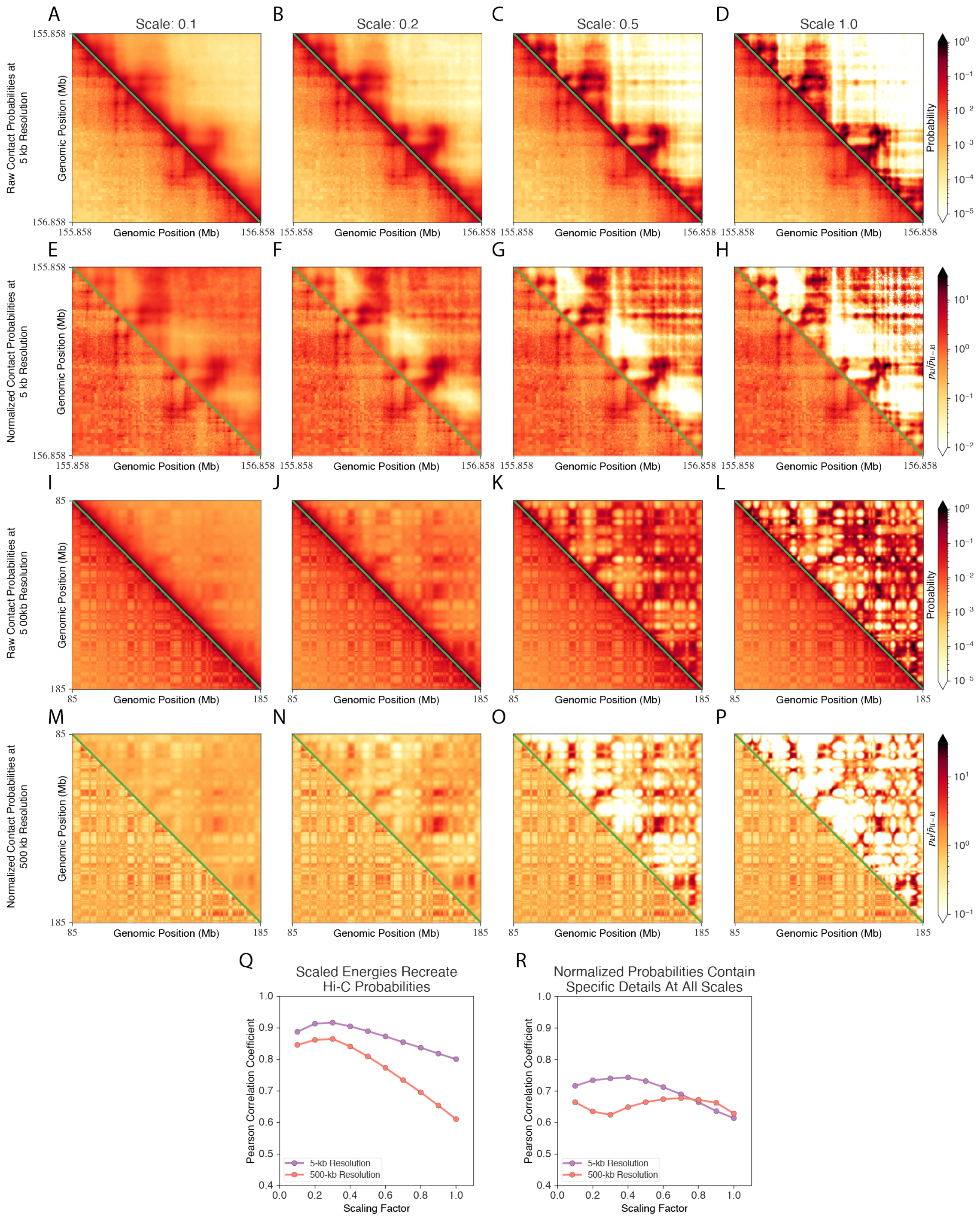


**Figure S11. The simulated contact statistics of polymer models parameterized by contact energies always contain local information, but the contact energies must be scaled to adequately exploring configurations.** (A-P) The simulated (upper triangle) and Hi-C (lower triangle) contact probabilities are compared in their raw form (A-D, I-L) and when normalized by genomic separation (E-H, M-P). Each row uses a consistent colormap. Contact probability maps display the simulated probabilities of polymer models parameterized by $\alpha_{kl} = 0.1\epsilon_i$ (A, E, I, M), $\alpha_{kl} = 0.2\epsilon_i$ (B, F, J, N), $\alpha_{kl} = 0.5\epsilon_i$ (C, G, K, O), or $\alpha_{kl} = 1.0\epsilon_i$ (D, H, L, P). (Q) When parameterizing polymer models with contact energies, decreasing their value increases the agreement between simulated and Hi-C contact probabilities. (R) Regardless of the scale applied to contact energies when parameterizing polymer models, the simulated probabilities contain similar information regarding local interactions.
